# Stress-induced clustering of the UPR sensor IRE1α is driven by disordered regions within its ER lumenal domain

**DOI:** 10.1101/2023.03.30.534746

**Authors:** Paulina Kettel, Laura Marosits, Elena Spinetti, Michael Rechberger, Philipp Radler, Isabell Niedermoser, Irmgard Fischer, Gijs A Versteeg, Martin Loose, Roberto Covino, G Elif Karagöz

## Abstract

Upon accumulation of unfolded proteins at the endoplasmic reticulum (ER), IRE1 activates the unfolded protein response (UPR) to restore protein-folding homeostasis. During ER stress, IRE1’s ER lumenal domain (LD) drives its clustering on the ER membrane to initiate signaling. How IRE1’s LD assembles into high-order oligomers remains largely unknown. By *in vitro* reconstitution experiments we show that human IRE1α LD forms dynamic biomolecular condensates. IRE1α LD condensates were stabilized when IRE1α LD was tethered to model membranes and upon binding of unfolded polypeptide ligands. Molecular dynamics simulations suggested that weak multivalent interactions are involved in IRE1α LD assemblies. Mutagenesis showed that disordered regions in IRE1α LD control its clustering *in vitro* and in cells. Importantly, dysregulated clustering led to defects in IRE1α signaling. Our results reveal that membranes and unfolded polypeptides act as scaffolds to assemble dynamic IRE1α condensates into stable, signaling competent clusters.

## Introduction

The endoplasmic reticulum (ER) controls various fundamental cellular functions ranging from folding and quality control of secreted and membrane proteins to lipid biogenesis. A set of conserved signaling pathways collectively known as the unfolded protein response (UPR) maintains ER homeostasis ^1^. IRE1, a single-pass ER transmembrane kinase/RNase, drives the most conserved UPR pathway ^2–6^. In response to ER stress, IRE1 assembles into clusters, which brings its cytosolic kinase and RNase domains in close proximity allowing for trans-autophosphorylation of the kinase domains and subsequent allosteric activation of its RNase domain ^7–10^. IRE1’s RNase activity initiates the nonconventional splicing of the mRNA encoding the transcription factor XBP1. The spliced form of *XBP1* mRNA drives expression of the genes involved in restoring ER homeostasis, including chaperones ^3, 4, 6, 10–15^. In metazoans, IRE1 activation also leads to the degradation of ER-bound mRNAs in a process known as regulated IRE1-dependent mRNA decay (RIDD), which decreases the ER protein-folding burden to alleviate ER stress ^16, 17^.

IRE1 senses various perturbations to ER homeostasis to initiate signaling. Under steady-state conditions, IRE1’s LD is bound by the ER chaperone BiP, which keeps IRE1 in an inactive state ^18^. Accumulation of misfolded proteins in the ER results in the dissociation of BiP from IRE1’s LD ^18–21^. Under these conditions, IRE1’s lumenal domain (LD) binds misfolded proteins as ligands that trigger its oligomerization ^22–24^. IRE1 can also sense lipid bilayer stress by its transmembrane domain leading to its activation ^25–27^. IRE1 activation highly correlates with its assembly into microscopically visible clusters in cells ^8, 22, 28–30^. Clustering of IRE1 is initiated by its ER-lumenal sensor domain ^7, 9, 22, 29, 31^. Importantly, mutations introduced to the oligomerization interface in IRE1’s LD impair the formation of high-order oligomers and abolish IRE1 signaling in cells ^7, 22, 29, 31^.

Oligomerization is a conserved property of IRE1 LD from yeast to humans ^7, 22^. *In vitro,* the core folded domain of human IRE1α LD (cLD) forms discrete dimers, which in a concentration-dependent manner assemble into dynamic high-order oligomers ^22^. The human IRE1α cLD was crystallized as a monomer in the unit cell and the crystal structure did not display functional oligomerization interfaces ^21^. Therefore, the structural basis for IRE1α LD oligomerization has remained elusive. Mutational analyses based on crosslinking coupled to mass spectroscopy data identified a hydrophobic segment in IRE1α cLD that controls its oligomerization *in vitro* and its clustering in cells ^22^. However, this method did not provide sufficient resolution to map the interfaces contributing to the formation of high-order oligomers. Therefore, the mechanistic basis of IRE1α oligomerization and the states leading to formation of signaling competent IRE1α oligomers have been poorly understood. Importantly, even though IRE1α is a membrane protein, how the two-dimensional physiological orientation of IRE1α on the membrane impacts its clustering has not been explored.

To mechanistically dissect how IRE1α LD assembles into high-order oligomers, we reconstituted IRE1α LD clustering in solution and on supported lipid bilayers (SLB) as model membranes. We revealed that disordered regions (DRs) in IRE1α LD control its assembly into dynamic biomolecular condensates. Our data suggest that membranes and unfolded polypeptide ligands act synergistically in stabilizing dynamic IRE1α LD condensates into long-lived clusters to transmit the signal across the ER membrane.

## Results

### IRE1α LD forms stable clusters on synthetic membranes

To investigate whether membrane association influences IRE1α LD clustering, we reconstituted the system *in vitro* using purified human IRE1α LD tethered to supported lipid bilayers (SLBs). SLBs are constituted of planar membranes formed on solid surfaces which are widely used as membrane-mimics (**Fig. 1A,B**).

**Fig. 1.**
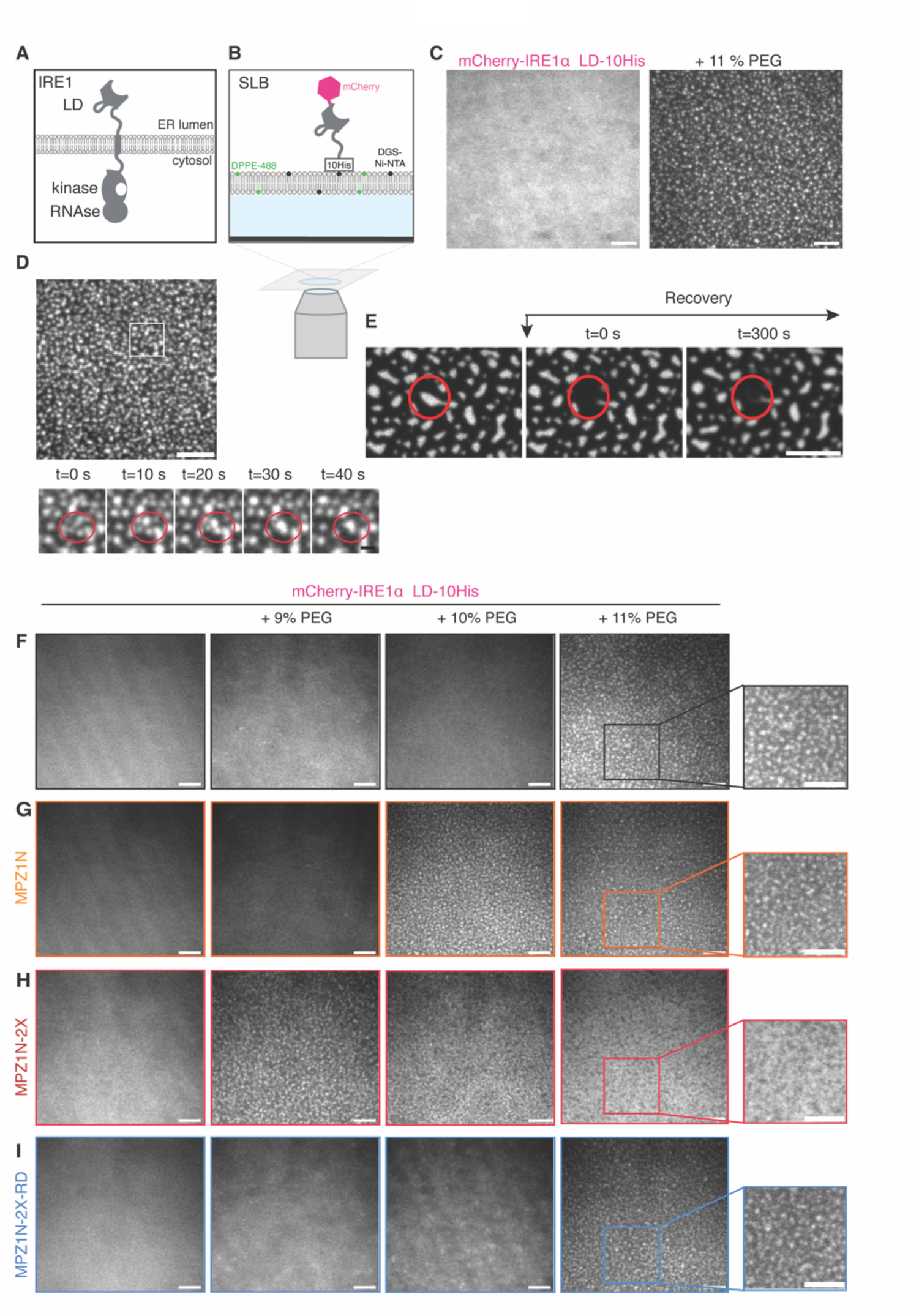
IRE1α LD forms clusters on supported lipid bilayers (SLB). **A.** Schematic illustration of IRE1α domain architecture within the ER membrane. **B.** Schematic illustration of the SLB setup. **C.** TIRF images of mCherry-IRE1α LD-10His clustering on an SLB in the absence (left) and presence of 11 % PEG. Scale bar (SB) = 5 µm. **D.** Fusion events of mCherry-IRE1α LD-10His clusters on SLBs at the indicated time points. Scale bar = 5 µm, zoom in scale bar = 1 µm. **D.** FRAP images of mCherry-IRE1α LD-10His on SLBs in presence of 11 % PEG within 300 s. Scale bar = 5 µm. **E.** TIRF images displaying clustering of mCherry-IRE1α LD-10His tethered to SLBs *via* 1 % Ni-NTA lipids in the presence of the indicated concentrations of PEG **F.** Clustering is visible by the formation of fluorescent intense spots. Scale bar = 5 µm **G.** TIRF images displaying clustering of mCherry-IRE1α LD-10His in the presence of PEG and 10 µM model unfolded polypeptide ligand MPZ1N. Scale bar = 5 µm. **H.** TIRF images displaying the phase diagram of mCherry-IRE1α LD-10His in the presence of PEG and 1 µM model unfolded polypeptide ligand MPZ1N-2X. Scale bar = 5 µm. **I.** TIRF images displaying the phase diagram of mCherry-IRE1α LD-10His in the presence of the indicated concentrations of PEG and 1 µM control peptide MPZ1N-2X-RD. Scale bar = 5 µm.

We reconstituted SLBs composed primarily of 1-palmitoyl-2-oleoyl-glycero-3-phosphocholine (98.92 mol% POPC). We used 1 mol% nickel–nitrilotriacetic acid (Ni-NTA) lipids to tether mCherry-IRE1α LD-10His to SLBs through its C-terminal 10xHis tag, which allows placing IRE1α LD in the topologically correct orientation (**Fig. 1B**). To monitor SLB integrity and fluidity, we used 0.08 mol% Atto488 labeled 1,2-Dipalmitoyl-sn-glycero-3-phosphoethanolamine (DPPE). Fluorescence recovery after photobleaching (FRAP) experiments revealed that mCherry-IRE1α LD-10His displayed a dynamic behavior on SLBs (**Fig 1C (left), Fig. Supp. 1A, Suppl. Table 1**). FRAP of Atto488 labeled DPPE lipids confirmed that the membrane was fluid (**Fig. Supp. 1A, Supp. Table 1**).

To mimic the crowding of the ER environment in our *in vitro* assays, we used the molecular-crowding agent polyethylene glycol 8000 (PEG) ^32^. We monitored mCherry-IRE1α LD-10His clustering *via* total internal reflection fluorescence (TIRF) microscopy and FRAP experiments at various PEG concentrations (**Fig. 1C (right), Fig. 1F, Fig. Supp. 1B,C**). Using the recovery half-life times obtained by the FRAP experiments, we calculated the diffusion coefficient based on Axelrod et al^33^ and Soumpasis et al^34^. Increasing the PEG concentration gradually decreased the mobile fraction and diffusion rates of mCherry-IRE1α LD-10His from 0.18 µm^2^/s without PEG to 0.02 µm^2^/s in presence of 11 % (w/v) PEG, demonstrating that the diffusion rate of IRE1α LD on SLBs decreases in the presence of crowding agent (**Fig. Supp. 1B-C**). In the presence of 10 % PEG, mCherry-IRE1α LD-10His displayed a diffuse fluorescence signal (**Fig. 1F, Fig. Supp. 1B, Supp. Table 2**), while 11 % PEG induced the formation of large mCherry-IRE1α LD-10His clusters on the SLB (**Fig. 1C (right), Fig. 1F, Fig. Supp. 1B,C, Movie 1, Supp. Table 2**). In the presence of 11 % PEG, both mCherry-10His control and Atto488-labeled DPPE retained their dynamic behavior confirming that the integrity of the SLB was not compromised and clustering is specific to IRE1α LD (**Fig. Supp. 1D-F, Supp. Table 2**). Under those conditions, mCherry-IRE1α LD-10His clusters formed and fused over time (**Fig. 1D, Movie 2**). Yet, FRAP experiments showed that photo-bleached IRE1α LD clusters did not recover even after 300 seconds (**Fig. 1E, Fig. Supp. 1B**). Instead, we observed a slight increase in mCherry-IRE1α LD-10His fluorescence at the periphery of the clusters (**Fig. 1E**) indicating that membrane-tethered IRE1α LD assembles into stable clusters driven by molecular crowding. To test whether membrane-tethered IRE1α LD clusters are not just aggregates, we performed wash-out experiments in which we removed the crowding agent from the well. Removal of PEG led to the disappearance of IRE1α LD clusters back to a diffuse fluorescence signal (**Fig. Supp. 1G**). Importantly, clusters could reform by adding 11 % PEG, indicating that they are dynamic and reversible. Altogether, we found that IRE1α LD forms stable but reversible clusters on synthetic membranes. Notably, our data are in line with the FRAP experiments performed with IRE1α in cells indicating that IRE1α LD reconstituted on membranes recapitulates the physical properties of IRE1α assemblies in cells ^30^.

### Binding of model unfolded polypeptides enhances IRE1α LD clustering

IRE1α’s LD binds unfolded peptides that are enriched in arginine, aromatic and hydrophobic residues as a means of recognizing aberrant protein conformations ^22, 23^. We next tested whether binding of model unfolded polypeptides would enhance IRE1α LD clustering on SLBs. We used peptides that we had previously shown to interact with IRE1α LD ^22^. The binding peptides with the highest affinity were derived from Myelin Protein Zero (MPZ) referred to as MPZ derivatives. MPZ1N is a 12mer peptide with a single binding site for IRE1α LD and binds IRE1α LD with an approximate affinity of 20 µM (**Fig. Supp. 2A**). MPZ1N-2X consists of two MPZ1N 12mers arranged in tandem, and it binds IRE1α LD with 1 µM affinity due to avidity ^22^ (**Fig. Supp. 2B**). As a control, we mutated arginine residues in MPZ1N-2X to impair its interaction with IRE1α LD, yielding MPZ1N-2X-RD ^22^. Using fluorescence anisotropy experiments, we confirmed the MPZ1N-2X-RD interaction with IRE1α LD is largely impaired (**Fig. Supp. 2B**).

In the stressed ER, IRE1α LD clustering may be initiated by specific interactions of IRE1α LD with un/misfolded proteins, and a bulk increase in molecular crowding due to blocked secretion of un/misfolded proteins. Therefore, we next tested whether IRE1α LD’s interactions with model unfolded polypeptides would decrease the threshold for its clustering in the presence of a crowding agent. We found that incubation with peptides reduced the effective concentration of PEG required to drive the clustering of mCherry-IRE1α LD-10His (**Fig. 1F-H, Fig. Supp. 2C**). This increased propensity was specific, as incubation of mCherry-IRE1α LD-10His with the mutant peptide MPZ1N-2X-RD did not impact its clustering (**Fig. 1I**). Importantly, the FRAP experiments revealed that the peptides did not impair SLB integrity (**Fig. 1F-I, Fig. Supp. 2D-F, Supp. Table 2**). In sum, we succeeded in reconstituting ligand-enhanced IRE1α LD clustering on synthetic membranes from minimal components, thus recapitulating a critical step of the UPR.

### IRE1α LD forms dynamic condensates in solution

Our data suggested that IRE1α LD tethered to synthetic membranes forms stable clusters due to restricted conformational freedom on planar surfaces. This model predicts that IRE1α LD clusters formed in solution should exhibit a more dynamic behavior when compared to those formed on SLBs. To test this prediction, we monitored IRE1α LD clustering in solution by differential interference contrast (DIC) microscopy. The systematic analyses of protein concentrations and buffer conditions by DIC showed that in the presence of 6 % PEG, 12,5 µM IRE1α LD formed droplets in solution. IRE1α LD droplets resembled biomolecular condensates formed through liquid-liquid phase separation (LLPS). Both the number and size of the condensates increased at higher protein concentrations (**Supp. Fig. 3A,B**) IRE1α LD condensates displayed dynamic and liquid-like behavior in solution, as evidenced by fusion events (**Fig. 2A,B, Movie 3**.). FRAP experiments confirmed the liquid-like nature of IRE1α LD condensates and revealed that IRE1α LD molecules exchanged in and out of the condensates with a mobile fraction of 88 % (t_1/2_ = 173.4 s); **Fig. 2C** and **Movie 2**). It has been observed that if a protein goes through LLPS, the liquid droplets will wet the glass surface, whereas hydrogels or less dynamic condensates do not wet solid surfaces or change shape ^35^. IRE1α LD condensates wetted the bottom of the glass surface in a time-dependent manner, confirming their liquid-like properties (**Supp. Fig. 3C**). Neither IRE1α LD^D123P^ mutant, which is impaired in dimerization ^21^, nor the mCherry control formed condensates. This suggested that D123 is required for formation of larger IRE1α LD clusters in solution (**Fig. Supp. 3D**). Altogether, our data revealed that in solution IRE1α LD forms dynamic condensates upon molecular crowding. These data suggested that tethering IRE1α LD to membranes leads to stabilization of IRE1α LD assemblies in the condensates. It is plausible that the restriction of IRE1α LD’s degree of freedom, or membrane-induced structural rearrangements, stabilize interfaces important for its clustering. Consequently, this could drive formation of long-lived IRE1α LD assemblies on membranes.

**Fig. 2.**
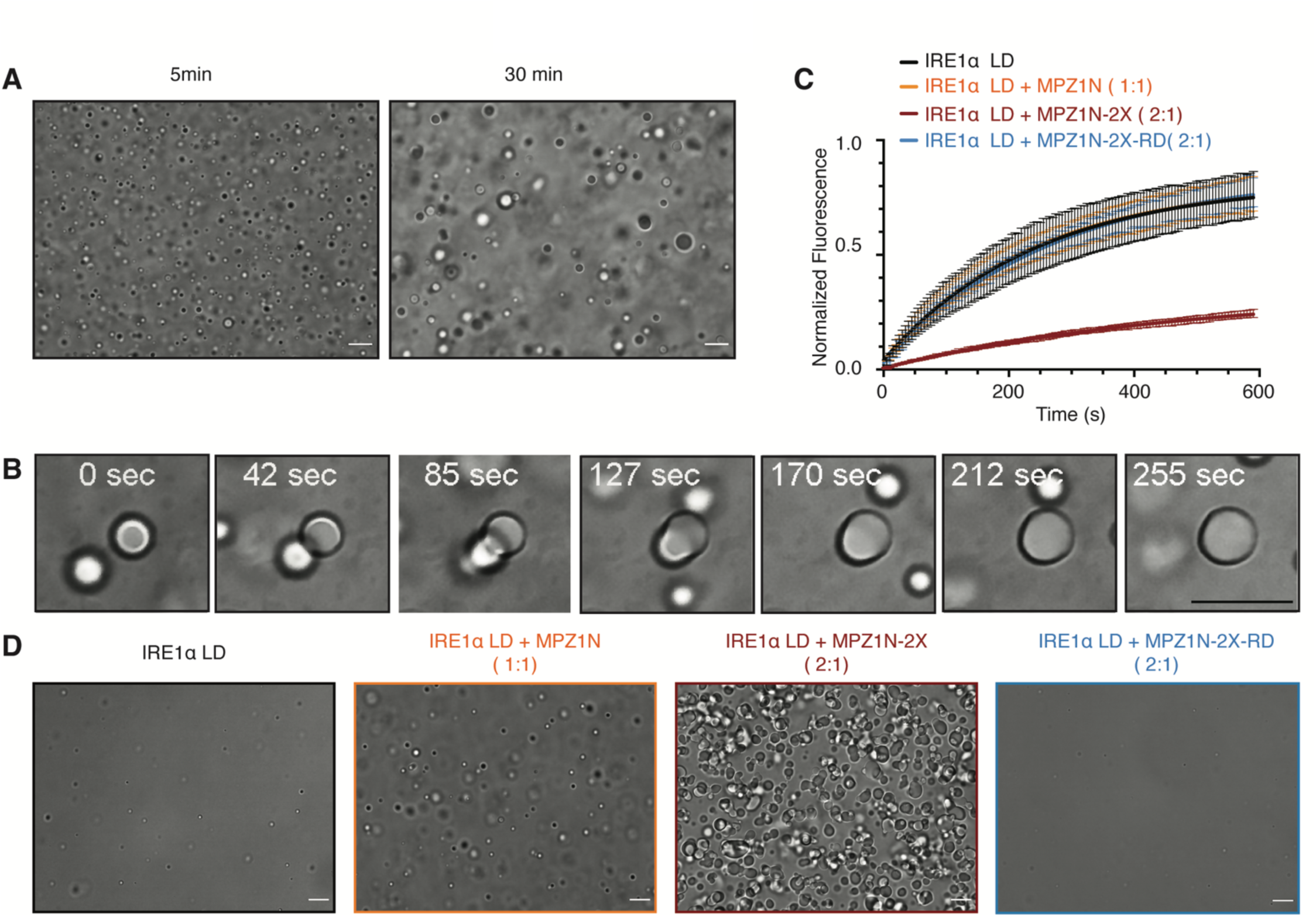
IRE1α LD forms dynamic condensates in solution. **A.** DIC microscopy images showing IRE1α LD condensates imaged after 5 min (left) and 30 min incubation with PEG (50 µM IRE1 LD, 6 % PEG). Scale bar for all images = 10 μm. **B.** Fusion of IRE1α LD condensates imaged by DIC microscopy. The condensates were imaged after 30 min incubation with PEG at the indicated time points (50 µM IRE1 LD, 6 % PEG). **C.** FRAP curves showing normalized fluorescent recovery of IRE1α LD condensates after 30 min incubation with 6 % PEG. IRE1α LD (25 µM IRE1 LD, 6 % PEG) in the absence (black curve) and in the presence of MPZ1N peptide (1:1 stoichiometry, orange curve), in the presence of MPZ1N-2X peptide (2:1 stoichiometry, dark red curve), in the presence of MPZ1N-2X-RD peptide (2:1 stoichiometry, blue curve). Curve marks show the mean value, error bars display the standard deviation. n=3 independent experiments were performed while 3 condensates were bleached each experiment. **D.** DIC microscopy images displaying LLPS behavior of IRE1α LD alone (50 µM IRE1α LD, 5% PEG) (left) and IRE1α LD in complex with MPZ1N (1:1 stoichiometry), MPZ1N-2X (2:1 stoichiometry) and the control MPZ1N-2X-RD. Images were taken 30 min after induction of phase separation with PEG.

We next characterized the impact of unfolded polypeptides on the formation and dynamics of IRE1α LD condensates. Fluorescein labeled MPZ1N-2X efficiently partitioned into preformed IRE1α LD condensates, revealing that they recruit client proteins (**Supp. Fig. 3E, left panel**). Instead, the Fluorescein-MPZ1N-2X-RD control peptide was not enriched in the condensates (**Supp. Fig. 3E, right panel**). In an experimental condition where IRE1α LD barely formed condensates (**Fig. 2D, left panel**), its incubation with stoichiometric amounts of model unfolded peptides led to the formation of large condensates (**Fig. 2D, Fig. Supp. 3F**). Instead, the control peptide MPZ1N-2X-RD did not impact IRE1α LD phase separation (**Fig. 2D, right panel, Fig. Supp. 3F**). Importantly, model unfolded polypeptides did not undergo phase separation in those conditions (**Supp. Fig. 3G,H**). This data indicated that specific interactions with unfolded polypeptides facilitate IRE1α LD phase separation. We next assessed whether unfolded polypeptide-binding would impact the dynamics of IRE1α LD assemblies in the condensates. FRAP experiments showed that while MPZ1N did not significantly impact IRE1α LD’s half-time recovery after photobleaching, instead binding of MPZ1N-2X peptide led to an increase in the recovery time of IRE1α LD (**Fig. 2C, Fig. Supp. 3I-J, Supp. Table 3**). These data revealed that a peptide with a single binding site shifts IRE1α LD to a confirmation that favors its clustering consistent with previous findings ^22^. MPZ1N-2X, which has two binding sites, can nucleate clusters by bridging IRE1α LD molecules and further stabilize IRE1α LD assemblies. We speculate that binding of unfolded polypeptides with various stoichiometry and biochemical properties might tune dynamics of IRE1α condensates and impact UPR signaling in cells.

### Disordered regions in IRE1α LD drive dynamic clustering

The biomolecular condensates formed by IRE1α LD in solution prompted us to ask which molecular interactions might explain this behavior. The formation of biomolecular condensates is often controlled by disordered regions in proteins ^36^. IRE1α LD comprises a mostly folded N-terminal motif (aa 24-307) joined to the transmembrane helix by a disordered region (aa 307-443) (**Fig. 3A, Fig. Supp. 4A**) ^37^. In the crystal structure of IRE1α cLD (aa 24-390, pdb: 2hz6 ^21^), several segments (i.e. (aa) 131-152, 307-358, and 369-390) are not resolved due to their flexibility (**Fig. 3B, Fig. Supp. 4A,B**). We refer to the disordered regions in IRE1α LD as Disordered Region 1 (DR1, aa 131-152), Disordered Region 2 (DR2, aa 307-358), Disordered Region 3, (DR3, aa 369-390), and the linker region (aa 391-443), respectively. Here, we employed molecular dynamics (MD) simulations to characterize their conformation and interaction.

**Fig. 3.**
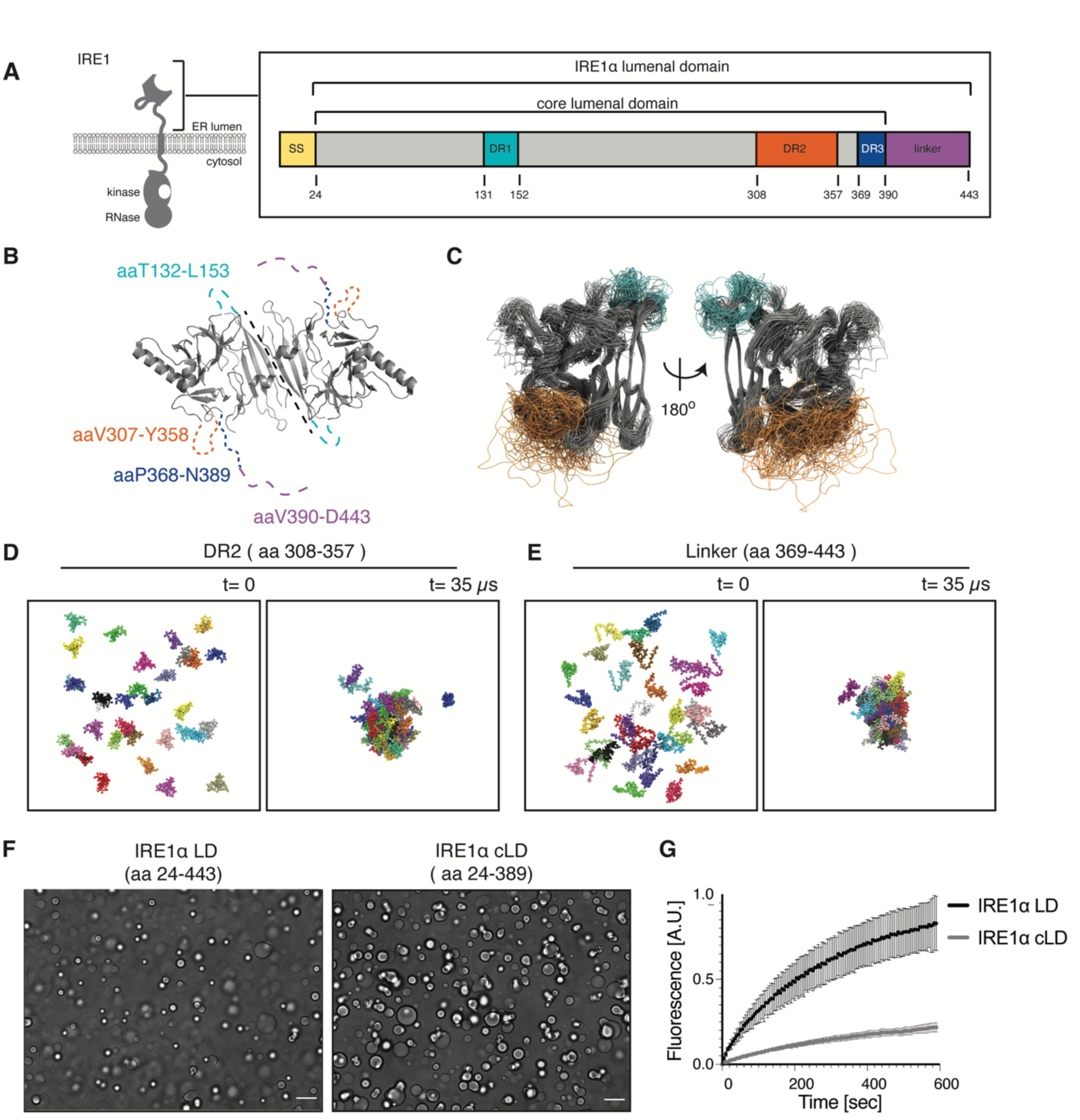
Disordered regions in IRE1α LD have potential to form clusters. **A.** Schematic description of DRs in IRE1α LD and boundaries of the core LD. The numbers correspond to the amino acid number at the domain boundaries. SS = Signal sequence, DR = disordered region. **B.** IRE1α cLD dimeric structure based on the crystal structure of human IRE1α (pbd: 2hz6). The DRs that are not resolved in the structure are depicted by dashed lines. **C.** Superposition of frames of an all-atom cLD simulation. Molecular dynamic simulations of IRE1α cLD shows flexibility of the DR at 600 ns time scale. **D.** Molecular Dynamics Simulations of 33 copies of DR2. These simulations reveal that the DR2 region forms clusters. **E.** Molecular Dynamics Simulations of 33 copies of the linker region. These simulations reveal that the linker region forms clusters. **F.** DIC images of IRE1α LD (left) and IRE1α cLD (right) reveal that IRE1α cLD is sufficient to form condensates. All images were obtained for 50 µM protein after incubation with 6 % PEG for 30 min. Scale bar = 10 μm. **G.** FRAP curve showing the time-dependent, normalized fluorescent recovery of 25 µM IRE1α LD and IRE1α cLD condensates after 30 min incubation with 6 % PEG. Curve marks show the mean value, error bars display the standard deviation and the values are fitted to a one-phase association curve displaying a lower mobile fraction and longer half-life time for IRE1α cLD condensates. n=3 independent experiments were performed where 3 condensates were bleached each experiment.

Atomistic MD simulations of the IRE1α cLD dimer (residues aa 29-368) revealed that DR1 and DR2 remain highly disordered during a 1 µs long simulation, not adopting any distinct secondary structures (**Fig. 3C**). DR2 was the most flexible part of the dimer. These data are in line with the published hydrogen-deuterium exchange experiments ^38^. We then performed coarse-grained MD simulations to test whether the disordered regions might self-associate (**Fig 3D,E**). We observed that DR1 did not form clusters in a 20 µs-long simulation (**Fig. Supp. 4C**). Instead, DR2 and the linker region readily clustered after 1 µs of simulation. The clusters were highly dynamic, and we observed reversible association of single polypeptide chains. To test the potential of heterologous associations, we simulated a system containing DR1 and DR2 and another system comprising DR2 and the linker region (**Fig. Supp. 4C,D**). DR2 clusters did not interact with the DR1 segments, which remained free in solution, while the DR2 and the linker formed well-mixed clusters. These data suggested that DR2 and the linker have the potential to form protein condensates.

We next investigated which specific interactions may drive the disordered regions to cluster. In protein condensates, the contacts formed inside a single polypeptide chain often resemble the ones formed across different polypeptide chains ^39^, indicating that the same interactions promote the internal and oligomeric organization. Therefore, we computed contact maps for interactions formed within single polypeptide chains and across different polypeptide chains in the clusters. Indeed, 1-D plots derived from the contact matrices, where we summed up the contributions from all possible interactions that a single residue forms in the simulations, confirmed this feature. Interactions within and between the DR2 and the linker region were mainly formed by the charged and aromatic residues (Asp, Lys, Phe) (**Fig. Supp. 5A-D**). In DR2, Asp328 and Lys349 formed the most probable contacts, suggesting an important role in cluster formation (**Fig. Supp. 5A-D**). The contact analysis showed that distinct regions in the disordered segments in IRE1α LD have the propensity to form low-affinity transient interactions driven by aromatic and charged residues. In summary, MD simulations revealed the biochemical potential of the disordered segments in IRE1α’s LD in driving its LLPS.

### IRE1α cLD forms rigid condensates

As MD simulations predicted that DR2 and the linker region form clusters in isolation, we next tested their LLPS potential through DIC microscopy with the purified constructs. We found that the core lumenal domain (cLD aa 24-389), which lacks the disordered linker region, formed condensates (**Fig. 3F, right**). These data revealed that the linker region is not necessary for formation of IRE1α LD condensates. IRE1α cLD rapidly formed condensates at lower protein and PEG concentrations compared to IRE1α LD **(Fig. Supp. 3A,B vs Fig. Supp. 6A,B**). DIC microscopy revealed that IRE1α cLD (aa 24-389) formed structures that resembled beads on a string (**Fig. 3F, right**). IRE1α cLD condensates accumulated on the glass slide without wetting the surface (**Fig. Supp. 6C**). FRAP experiments showed that IRE1α cLD recovered after t_1/2_ = 281.5 s and displayed a 26.7 % mobile fraction confirming that cLD condensates are less dynamic in comparison to IRE1α LD condensates (**Fig. 3G, Supp. Table 4**). In support of the low mobile fraction of IRE1α cLD revealed by the FRAP data, mCherry-tagged IRE1α LD partitioned into preformed IRE1α LD condensates after 5 min, in contrast mCherry tagged-cLD failed to do so (**Fig. Supp. 6D**). Altogether, our data showed that IRE1α cLD formed stable condensates, indicating that the linker segment (aa 390-443) modulates both the propensity to coalesce and IRE1α LD associations in condensates. Together with the MD simulations, these data suggest that the linker region forms transient intra- and intermolecular contacts that potentially regulate a critical switch in UPR signaling.

### Mutations in IRE1α LD’s disordered regions modulate clustering *in vitro*

We next screened for mutants in the disordered segments that may regulate IRE1α LD’s clustering. For these experiments, we used LLPS assays in solution to rapidly screen for mutants that impair IRE1α LD self-assembly. As IRE1α cLD could readily form condensates in solution, we mutated DR2 and DR3 in IRE1α LD. We chose regions enriched in hydrophobic or aromatic sequences, which might form intermolecular contacts to nucleate phase separation ^40^. Specifically, we mutated 3- or 4-residue stretches to glycine-serine residues, which often form dynamic segments acting as spacers in biomolecular condensates (**Fig. 4A, Supp. Fig 4A**).

**Fig. 4.**
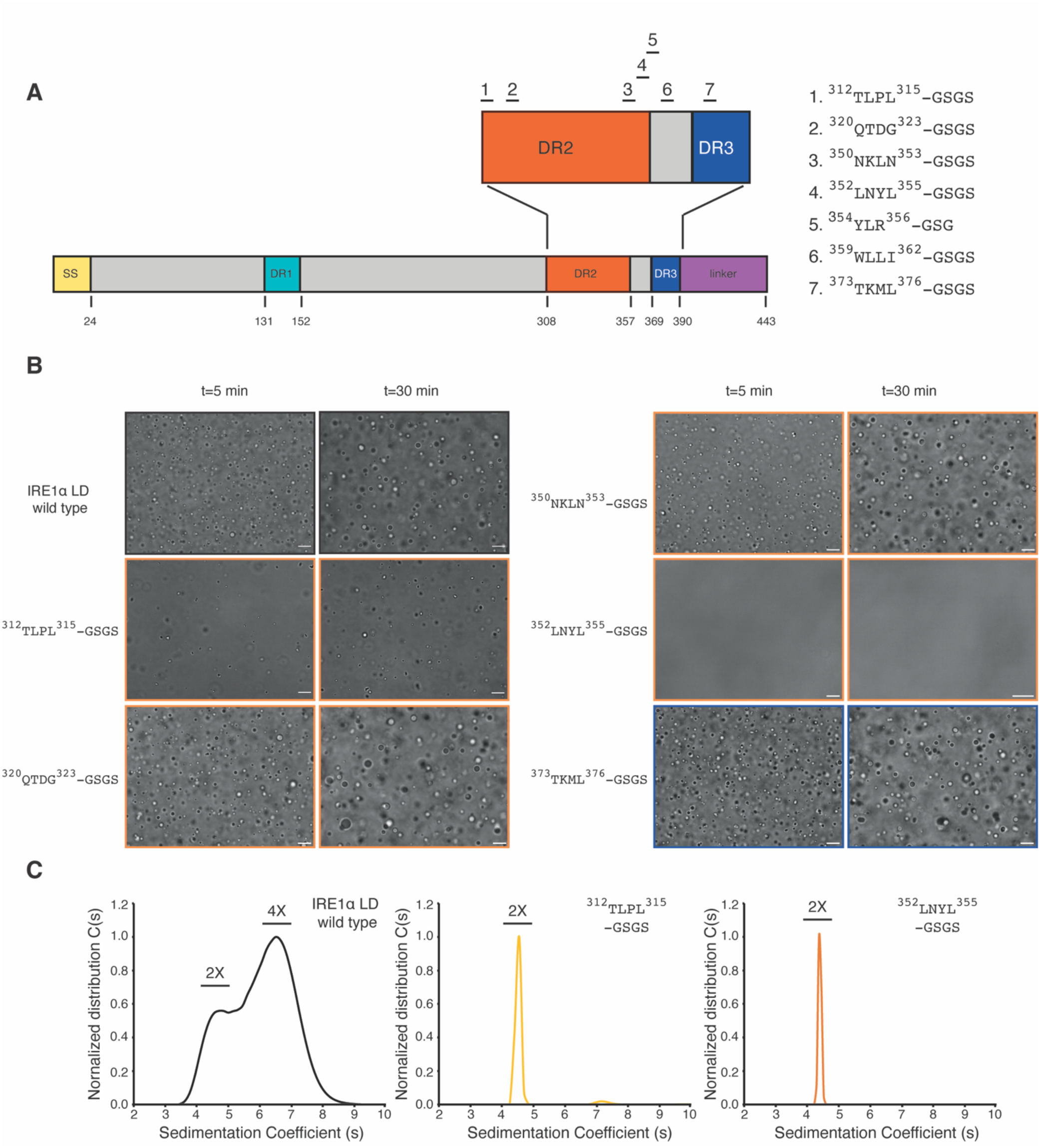
Mutations in disordered segments in IRE1α LD impair phase separation. Schematic description of the mutations (1-7) introduced to IRE1α LD. SS = Signal sequence, DR = disordered region. **B.** DIC images showing LLPS behavior of IRE1α LD wild type and the mutants at 50 µM after their incubation with 6 % PEG for 5 min (left) and 30 min (right). Scale bar = 10 μm. **C.** AUC-SV curves of 25 µM wild type IRE1α LD (black curve, left), IRE1α LD ^312^TLPL^315-^GSGS (light orange curve, middle) and IRE1α LD ^352^LNYL^355-^GSGS mutants (orange curve, right).

All the mutants were biochemically stable upon purification, allowing us to study their clustering behavior by DIC microscopy. The IRE1α LD^312^TLPL^315^ mutant formed smaller condensates with slower kinetics that failed to fuse efficiently suggesting that this region is important for condensate formation. A mutation in the ^320^QTDG^323^ segment did not impair phase separation (**Fig. 4B, Fig. Supp. 7A**), whereas mutating a segment enriched in hydrophobic and aromatic residues (IRE1α LD^352^LNYL^355^) abolished formation of IRE1α LD condensates (**Fig. 4B**). Introducing a mutation to the neighboring segment ^354^YLR^356^ strongly impaired LLPS (**Fig. Supp. 7B**), yet mutating the segment preceding it (^350^NKLN^353^) only slightly impacted LLPS (**Fig. 4B**). Similarly, the IRE1α LD ^373^TKML^376^ mutant in DR3 did not impact condensate formation (**Fig. 4B**). These data revealed that ^354^YL^355^ region forms a hot spot for molecular interactions driving IRE1α LD clustering.

Notably, ^354^YL^355^ resides near ^359^WLLI^362^, whose mutation to GSGS impairs IRE1α LD oligomerization ^22, 31^. DIC microscopy indicated that IRE1α LD ^359^WLLI^362^ mutant underwent LLPS, even in conditions in which wild-type IRE1α LD barely formed condensates (**Fig. Supp. 7C**). These results revealed that disrupting the canonical oligomerization interface did not hinder phase separation of IRE1α LD and, moreover, suggested that oligomers are distinct from condensates (**Fig. 4B, Fig. Supp. 7C**). These results motivated us to interrogate the oligomerization behavior of IRE1α LD ^352^LNYL^355^ and ^312^TLPL^315^ mutants using orthogonal methods. To this end, we performed analytical ultracentrifugation sedimentation velocity (AUC-SV) experiments to determine whether IRE1α LD mutants could form high-order oligomers. These experiments revealed that, similar to previous observations on IRE1α cLD ^22^, IRE1α LD was found in equilibrium of dimers and oligomers at 25 µM. Strikingly, under those conditions, IRE1α LD ^352^LNYL^355^ and ^312^TLPL^315^ mutants only formed dimers. These data revealed that ^352^LNYL^355^ and ^312^TLPL^315^ regions are important for the formation of high-order IRE1α LD oligomers (**Fig. 4C**). Altogether, our data converge on a model in which IRE1α LD dimers interact with each other in various conformations. These interactions are facilitated by the propensity of DRs to assemble into stable oligomers with a distinct preferred structure and into condensates with low affinity contacts with no fixed valence. We propose that the transient low affinity interactions are crucial in bringing IRE1α molecules in close proximity to drive formation of active IRE1α assemblies.

### The disordered regions in IRE1α LD are important for its clustering in cells

IRE1α forms dynamic clusters in cells experiencing ER stress ^28–30, 41^. To validate the role of disordered segments in IRE1α LD clustering, we established stable cell lines expressing wild type human IRE1α or IRE1α harboring mutants with impaired (IRE1α LD LNYL and TLPL mutants) or enhanced (IRE1α cLD Δlinker) its clustering, as determined by our *in vitro* assays (**Supp. Fig. 6A, Fig. 4B**). We introduced doxycycline-inducible transgenes encoding mNeonGreen (mNG) tagged variants of IRE1α into mouse embryonic fibroblasts (MEFs) deficient for both isoforms of IRE1 (IRE1 α^−/−^ and IRE1β^−/−^), and monitored IRE1α clustering by fluorescence microscopy. We introduced the mNG tag into IRE1α’s cytoplasmic flexible linker ^22, 28, 30^. In the absence of doxycycline, cells expressed low levels of IRE1α due to the inherent leakiness of the doxycycline-inducible system (**Fig. Supp. Fig 8A-D**). Under these conditions, the expression levels of IRE1α-mNG and its mutant variants were similar to the level of endogenous IRE1α observed in wild-type MEFs as assessed by Western blot analysis (**Fig. Supp. 8D**). When we treated the cells with the ER stress inducing drug tunicamycin, cells carrying IRE1α-mNG showed a modest reduction in *XBP1* mRNA splicing activity compared with wild type control MEFs, suggesting that the mNG-tag slightly impairs its activity (**Fig. Supp. 8E**).

The size of IRE1α clusters in cell depends on the protein concentration ^28, 31, 42^. While clusters formed at endogenous levels of IRE1α in most tissues are generally too small to overcome the diffraction limit of light, IRE1α forms microscopically visible clusters when it is ectopically expressed to levels 2-20 times over endogenous protein levels ^22, 28–31, 41, 42^. To visualize IRE1α clustering in mammalian cells with confocal microscopy, we overexpressed IRE1α in MEFs. IRE1α expression levels increased linearly with doxycycline concentration in the range of 25 to 400 nM (**Fig. Supp. 8C**). Treatment with 400 nM doxycycline led to expression levels roughly 30-fold over endogenous IRE1α in wild type MEFs (**Fig. Supp. 8F**). Confocal microscopy experiments in which we monitored IRE1α cluster formation indicated that at 400 nM doxycycline, but not 100 nM, cells expressing wild type IRE1α-mNG formed microscopically visible clusters in a stress-dependent manner (**Fig 5A, Fig Supp. 8G**).

**Fig. 5.**
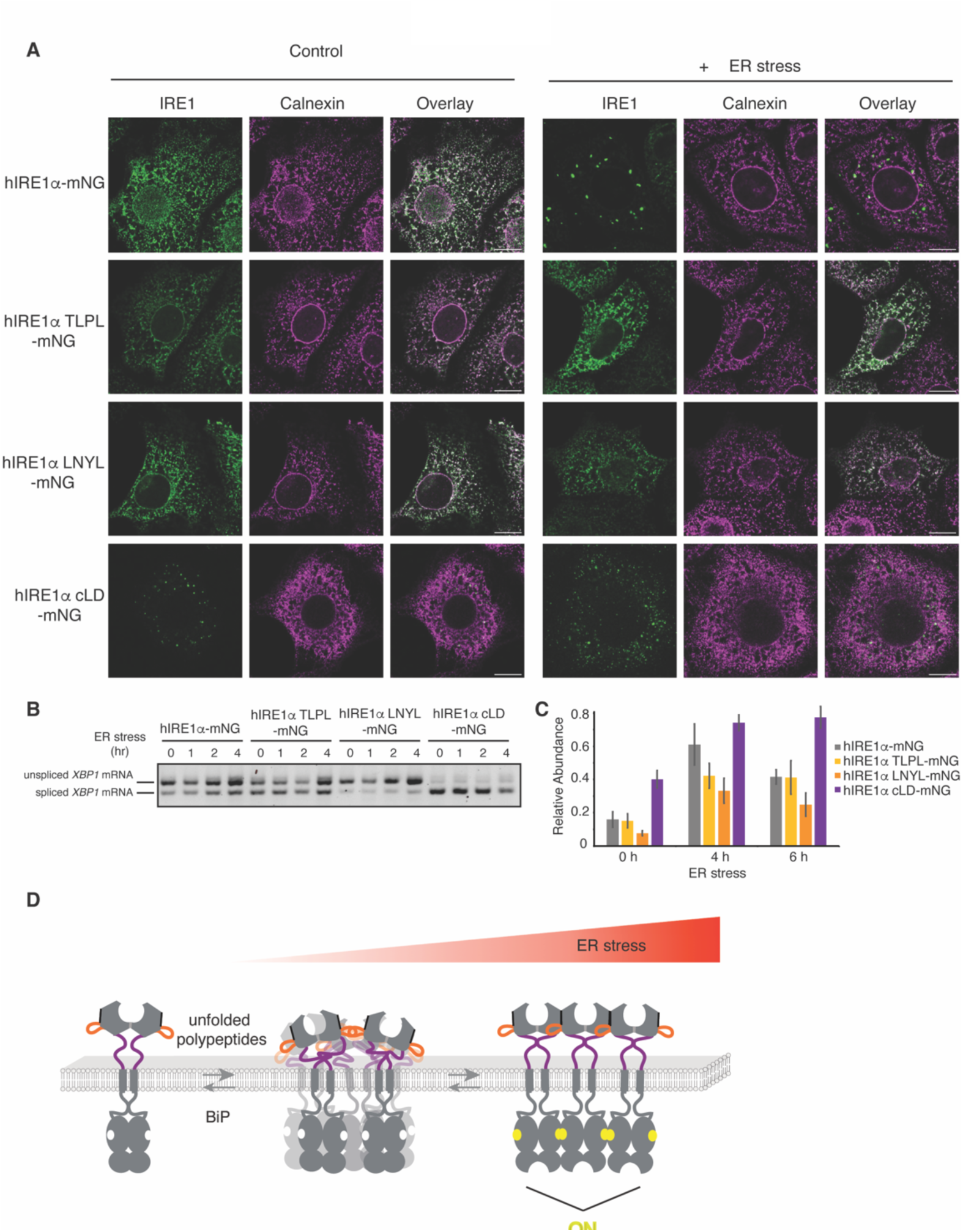
IRE1α LD DR mutants dysregulate its clustering and activity *in vivo*. **A.** Immunofluorescence images of MEFs treated with 400 nM doxycycline expressing IRE1α-mNG or its mutants in the absence (left panel) of stress and treated with 5 µg/ml ER stressor Tunicamycin for 4 hrs (right panel). IRE1α-mNG and its mutants are visualized by mNG fluorescence (green) and the ER-chaperone Calnexin is stained by anti-calnexin antibody (purple). Scale bar = 10 µm. **B.** Semiquantitative PCR reaction to monitor splicing of *XBP1* mRNA by IRE1α-mNG and its mutants at different time points after induction of ER stress by addition of 5 µg/ml Tunicamycin. Expression of the IRE1α variants is induced by treatment of MEFs 24 hrs with 400 nM doxycycline before induction of ER stress. The bands are indicated as unspliced and spiced *XBP1* variants. **C.** qRT-PCR to monitor splicing of *XBP1* mRNA by IRE1α-mNG and its mutants at different time points after induction of ER stress by addition of 5 µg/ml Tunicamycin. **D.** Model describing the role of DRs in IRE1α clustering. During ER stress, ER-resident chaperone BiP is released from the DRs in IRE1α LD allowing these segments to self-associate through multivalent weak interactions. Under those conditions, misfolded proteins accumulating in the ER facilitate formation of dynamic IRE1α condensates. These dynamic condensates rapidly assemble into stable IRE1α clusters with distinct conformation allowing for IRE1α trans-autophoshorylation and RNase activity.

Notably, in these conditions, IRE1α-TLPL-mNG and IRE1α-LNYL-mNG mutants failed to form visible clusters at comparable expression levels as the wild type IRE1α-mNG (**Fig 5A, Fig. Supp. 8A-F**). These results substantiated our findings that TLPL and LNYL segments in the LD are important for IRE1α assembly. In stark contrast, IRE1α cLD-mNG (Δlinker), which lacks the linker region, formed clusters constitutively in the absence of stress (**Fig 5A, Fig. Supp. 8G**), or even upon induction of its expression with a lower (100 nM) doxycycline concentration. This suggested that the clustering threshold is lower for IRE1α cLD-mNG consistent with the results we obtained in our *in vitro* experiments. Taken together, these results substantiate the role of disordered segments in IRE1α’s LD in regulating IRE1α’s self-association into high-order assemblies.

Next, we investigated whether the mutants would impact IRE1α activity monitored by its ability to splice *XBP1* mRNA. Semi-quantitative polymerase chain reaction (PCR) and quantitative real-time PCR (qRT-PCR) analyses revealed that IRE1α LNYL-mNG mutant exhibited impaired *XBP1* mRNA splicing activity compared to wild type IRE1α-mNG, indicating that the interfaces formed by the LNYL region in IRE1α LD play an important role in forming active IRE1α assemblies in cells (**Fig 5B,C**). Instead, IRE1α TLPL-mNG splicing activity was only slightly diminished ^31, 42^ (**Fig 5B,C**). These data suggested that the mutation in the interface formed by TLPL segment can be compensated by formation of other protein interfaces in cells. IRE1α cLD-mNG, which constitutively formed foci in the absence of stress, displayed high constitutive *XBP1* mRNA splicing activity **(Fig 5B,C**). These data revealed that the splicing activity of the mutants in cells is in excellent agreement with their ability to form biomolecular condensates *in vitro*. Altogether, these results indicate that disordered segments in the IRE1α LD are involved in the formation of signaling competent IRE1α assemblies in cells.

## Discussion

IRE1 governs the most evolutionarily conserved branch of the UPR. IRE1 signaling is tied to the formation of dynamic clusters in yeast and mammalian cells, and mutations that impair IRE1 clustering result in severely reduced activity ^8, 22, 28, 30^. Thus, self-assembly emerges as a fundamental principle of IRE1 regulatory control. IRE1 clustering is driven by its ER-sensor lumenal domain, which juxtaposes its cytosolic domains to activate its RNase domain. The structural features enabling IRE1α LD clustering and its mechanistic principles have remained unknown, and here, through bottom-up approaches to reconstitute IRE1α LD clustering in solution and on model membranes, we provide evidence for the role of DRs in regulating IRE1α’s self-assembly.

We found that the stress sensing LD of IRE1α formed dynamic biomolecular condensates in solution. In contrast, IRE1α LD formed long-lived clusters on model membranes similar to what was shown for IRE1α clusters in cells indicating that membrane-tethering stabilizes interactions among IRE1α LD molecules (**Fig. 1. C-E**), ^30^. We anticipate that these long-lived interactions are crucial for providing sufficient time for transmitting the information through the membrane bilayer to initiate the auto-phosphorylation of the kinase domains leading to activation of its RNase domain. This model is in line with the recent data, which showed a lag between IRE1α oligomerization and its trans-autophosphorylation activity in cells ^31^. In our experiments, we used a simple membrane composition, and thus future studies are necessary to assess how changes in the membrane composition during ER stress might regulate IRE1α clustering ^25, 26, 43^.

Biomolecular condensates are formed through multivalent low affinity interactions by the disordered segments in proteins ^44–47^. IRE1α LD has several DRs whose function has remained largely unknown. Surprisingly, the distinct DRs in IRE1α LD regulate the formation and dynamics of IRE1α LD clusters in opposite ways. Removing the linker region (aa 390-443), which connects the folded core domain to the transmembrane helix, decreased the clustering threshold of IRE1α. By contrast, mutating aromatic and hydrophobic amino acids in two distinct parts in the DR2 segment (^312^TLPL^315^ and ^352^LNYL^356^ mutants) impaired its oligomerization *in vitro*, and abolished formation of microscopically detectable clusters in cells.

One IRE1α LD mutant, IRE1α LD ^359^WLLI^362^, does not form oligomers^22^ but could readily form condensates even at lower protein concentrations. These data indicate that the distinct oligomeric conformation formed through the contacts provided by the ^359^WLLI^362^ segment is not required for the multivalent-transient interactions formed by ^312^TLPL^315^ and ^352^LNYL^356^ regions. Importantly, both ^359^WLLI^362^ and ^352^LNYL^356^ mutants display impaired *XBP1* mRNA splicing activity in mammalian cells indicating that they both contribute to the assembly of IRE1α into enzymatically active clusters ^22^. Our data suggest that these interfaces contribute to the formation of temporally separated distinct assembly intermediates to generate signaling competent IRE1α oligomers in cells. Intriguingly, recent correlated light and electron microscopy combined with electron cryo-tomography (cryo-CLEM-ET) imaging of the IRE1α clusters in mammalian cells suggested that IRE1α LD forms ordered double-helical filaments in its native, membrane-embedded state under stress conditions ^41^. We anticipate that the increased local concentration of IRE1α in the condensates might facilitate assembly of IRE1α LD into filaments with distinct structure observed in cells. The regions in IRE1α LD, which we identified to regulate its clustering, were earlier proposed to be recognized by the ER-chaperone BiP ^38^, and therefore, it is plausible that these regions are occluded by BiP binding, which could prevent clustering under non-stress conditions.

Clustering of IRE1α LD on membranes followed a sharp transition as a function of molecular crowding (compare 10 % and 11 % PEG, **Fig. 1F**), suggesting that an increase in ER protein load, as during ER stress, could constitute the sensing threshold for IRE1α ^48–53^. In line with our previous observations, we found that in addition to molecular crowding, IRE1α LD’s direct interaction with unfolded peptide ligands decreased the threshold for IRE1α LD clustering (**Fig. 1F-H**) and stabilized IRE1α LD condensates. We anticipate that misfolded proteins with diverse biochemical properties could differently modulate the threshold for IRE1α clustering and the stability of IRE1α clusters regulating both sensitivity and duration of the UPR in cells.

Our data converge on a model in which ER stress triggers BiP release from the DRs in IRE1α’s LD, allowing their association with DRs of other IRE1α molecules through low affinity transient contacts. ER stress increases molecular crowding in the ER due to secretory and protein-folding impairment and accumulation of misfolded proteins, both of which facilitate self-assembly of IRE1α LDs. Under those conditions, unfolded polypeptides that bind to IRE1α LD and membrane-imposed constraints further stabilize IRE1α clusters leading to the formation of stable IRE1α assemblies competent in UPR signaling (**Fig. 5D**).

LLPS of membrane-associated proteins has emerged as a novel mechanism regulating cellular organization and signaling ^54–58, 59^. Our data suggest that LLPS of IRE1α contributes to UPR signaling. IRE1α levels are controlled *via* intricate feedback loops that regulate protein abundance during ER stress ^60^ and aberrant overexpression of IRE1α in multiple myeloma and breast cancer contributes to pathology ^61, 62^. We anticipate that the novel assembly states of IRE1α identified here could be targeted by small molecules for therapeutic purposes in disease.

## Acknowledgements

We thank Thomas Peterbauer at the Max Perutz Labs Biooptics Light Microscopy Facility for his help and support. We are grateful to Kitti Csalyi and Thomas Sauer at Max Perutz Labs Biooptics FACS facility for their help. We acknowledge funding from Austrian Science Fund (FWF-SFB F79 and FWF-W 1261) to GEK. PK acknowledges the support of the Max Perutz PhD fellowship. We thank the members of the Karagöz lab for the critical reading and editing of the manuscript. We are thankful to our colleagues Diego Acosta-Alvear, Vladislav Belyy, Jirka Peschek, Yasin Dagdas, Javier Martinez, Sascha Martens and Alwin Köhler for their invaluable input on the manuscript. We are grateful to Life Science Editors, especially Katrina Woolcock for valuable input on the manuscript. GAV is funded by Stand-Alone grants (P30231-B, P30415-B, P36572), Special Research Grant (SFB grant F79), and Doctoral School grant (DK grant W1261) from the Austrian Science Fund (FWF). E.S. and R.C. acknowledge support and funding by the Frankfurt Institute of Advanced Studies, the LOEWE Center for Multiscale Modelling in Life Sciences of the state of Hesse, the Collaborative Research Center 1507 “Membrane-associated Protein Assemblies, Machineries, and Supercomplexes” (Project-ID Project ID 450648163), and the International Max Planck Research School on Cellular Biophysics (to R.C.), the Center for Scientific Computing of the Goethe University and the Jülich Supercomputing Centre for computational resources and support. We are thankful to Monika Kubickova for the help with the AUC experiments. We acknowledge CF BIC of CIISB, Instruct-CZ Centre, supported by MEYS CR (LM2023042)) and European Regional Development Fund-Project „UP CIISB“ (No. CZ.02.1.01/0.0/0.0/18_046/0015974).

## Movies

**Movie 1.** mCherry-IRE1α LD-10His cluster formation on SLBs after the addition of 11% PEG. Each frame is recorded every 2 sec for a total of 32 frames.

**Movie 2.** Fusion of mCherry-IRE1α LD-10His clusters on SLBs. The movie is recorded 10 min after induction of cluster formation by addition of 11 % PEG. Each frame is recorded every 2 sec for a total of 60 frames.

**Movie 3.** LLPS of IRE1α LD in solution. The movie is recorded 30 min after induction of LLPS with 6% PEG.

## Supplementary Tables

**Supplementary Table 1.** mCherry-IRE1 LD-10His Fluorescence Intensity on SLBs

**Supplementary Table 2.** Fits of the FRAP curves obtained from SLB tethered mCherry-IRE1 LD-10His and Atto488-DPPE.

**Supplementary Table 3.** Fits of the FRAP curves of mCherry-IRE1 LD-10His in condensates formed in solution.

**Supplementary Table 4.** Fits of the FRAP curves of mCherry-IRE1 LD-10His or mCherry-IRE1 cLD-10His in condensates formed in solution.

## Materials and Methods

### Generation of Constructs for in-vitro Assays

All constructs were constructed in pET-47 b(+) vector. hIRE1α LD mutants ^312^TLPL^315^-GSGS, ^350^NKLN^353^-GSGS, and ^352^LNYL^355^-GSGS were based on the T274C variant of hIRE1α LD (Cysteines are substituted by Alanins, Threonine aa274 was substituted by Cysteine). All other mutants were based on the WT hIRE1α LD. We could not observe any differences between hIRE1α LD WT and T274C in our assays. ^320^QTDG^323^-GSGS and ^373^TKML^376^-GSGS were constructed through site-directed mutagenesis, with subsequent blunt end ligation. For mCherry tagged proteins, an N-terminal mCherry sequence and C-terminal 10HisTag was used.

### Protein expression and purification

hIRE1α LD expression and purification was adapted from published protocols ^22^. In brief, *Escherichia coli* strain BL21DE3* RIPL was grown with the respective antibiotics in Luria Broth at 37°C until OD_600_ = 0.6-0.8. The protein expression was induced with 400 µM IPTG for hIRE1α variants without and 1 mM for variants with mCherry at 20°C and grew overnight. Before lysis and after each purification step, 1X Roche cOmplete Protease Inhibitor Cocktail was added to the cells or fractions containing protein. Cells were harvested and lysed (50mM HEPES pH 7.2-pH 7.4, 400 mM NaCl, 20 mM Imidazol, 5 mM β-mercaptoethanol, 0 or 10 % Glycerol) in an Avestin EmulsiFlex-C3 cell disruptor at 16,000 psi. The lysate was spun at 30,700 x g for 45 min. The supernatant was applied to a 5 ml His-TRAP column (GE Healthcare) and eluted with a gradient of 20 mM to 500 mM imidazole. The eluate was diluted with 50 mM Hepes pH 7.2 – pH 7.4 (in the absence or in the presence of 10 % Glycerol, 5 mM β-mercaptoethanol) to a concentration of 50 mM NaCl, to apply it to a HiTRAP Q HP (5 ml, GE Healthcare) anion exchange column. The protein was eluted with a linear gradient from 50 mM to 1 M NaCl. To remove the His tag from hIRE1α LD without mCherry, the protein was incubated with 3C Precision protease at a ratio of 50 to 1 over night at 4°C. The protein was loaded to a His-TRAP column before it was further purified on a Superdex 200 10/300 gel filtration column (25/50 mM HEPES pH 7.2 – pH 7.4, 150 mM NaCl, 5 mM DTT; for mCherry proteins: 25 mM HEPES pH 7.4, 150 mM KCl, 10 mM MgCl_2_, 5 mM DTT). The Expasy ProtParam tool (http://web.expasy.org/protparam/) was used to determine the extinction coefficient at 280 nm to get the final protein concentration.

### SLB preparation and assays

The protocol was adapted from (Bakalar et al. 2018, Cell^63^).

#### SUV preparation

In brief, SUVs were prepared by creating a dried lipid film of mainly POPC (1-palmitoyl-2-oleoyl-sn-glycero-3-phosphocholine, Avanti), 1 mol% Ni-NTA (DGS-NTA(Ni) (1,2-dioleoyl-sn-glycero-3-[(N-(5-amino-1-carboxypentyl)iminodiacetic acid)succinyl] (nickel salt)), Avanti) and 0.08 mol% Atto488 labeled DPPE (1,2-Dipalmitoyl-sn-glycero-3-phosphoethanolamine labeled with Atto488, Sigma-Aldrich) with an argon stream followed by desiccation for 45 min. The rehydration was performed with deionized water by gently vortexing followed by 40 s tip sonication at 20 % power for three times with 20 s in between to prevent generation of heat. The SUVs were filtered through a 0.22 µm PES filter (Carl Roth) and stored at 4°C for a maximum of 48 hrs to prevent oxidation of lipids.

#### SLB preparation

SLBs were formed in a silicone chamber (Grace Bio-Labs, GBL103280) sealed on an RCA cleaned (Nguyen *et al.* (2015) Methods Cell Biol.) 1.5 H, 24 x 50 mm coverslip (Carl Roth) by fusing 20 µl SUVs with 30 µl MOPS buffer (25 mM MOPS pH 7.4, 125 mM NaCl) for 10min at room temperature. The SLB was washed with 50 µl PBS and 50µl wash buffer (25 mM Hepes, pH 7.3, 150 mM NaCl, 250 µM TCEP) each four times, respectively. To determine the optimal concentration of protein in our reconstituted system, we incubated SLBs with various concentrations of mCherry-IRE1α LD-10His ranging from 50 nM to 500 nM, and set on a concentration of 200 nM, which was below saturation of the 1 mol% Ni-NTA lipids as determined by fluorescent intensity as a function of protein concentration (Supp. Table 1).

The protein was attached at a concentration of 200 nM by incubating for 10 min followed by three more wash steps with wash buffer to remove any unattached protein from the solution. Imaging was conducted on an Olympus cellSens Live Imaging TIRF system with an Olympus 100 × 1.49 NA high-performance TIRF objective with 7 % 488, 100 ms exposure and 10 % 561, 100 ms exposure *via* a Hamamatsu ImagEM X2 EM-CCD camera operated by Olympus cellSens 3.1.1. The fluidity of the membrane was confirmed *via* FRAP experiments. A 2 s 50 % single-point laser pulse of 405 nm was used to bleach the fluorescence of the membrane and protein and the fluorescence recovery was followed over 100 frames every 2 s. Image processing was performed in ImageJ by selecting the FRAP ROI and another ROI of the same size on a non-FRAPed area as bleaching background and was kept the same within an experiment. The bleaching ROI was used to obtain bleaching factors by which the FRAP values were corrected with, followed by normalization. The normalized FRAP values of all 100 frames for mCherry proteins and 15 frames for Atto488 labeled DPPE was fitted to an exponential recovery with no offset curve in ImageJ. The half-life time was used to calculate the diffusion coefficient based on Axelrod et al. and Soumpasis et al ^33, 34^ assuming only 2D diffusion of the protein on the SLBs as any access unbound protein was washed out. Image processing was performed in ImageJ adjusting the brightness and contrast of the images to be the same within a Figure panel.

#### Crowding assay in 2D

The protein of interest was incubated with the desired PEG concentration in 25 mM Hepes pH 7.3, 150 mM NaCl, 250 µM TCEP and PEG (indicated in the Figure legend) for 10 min before FRAP experiments were performed to access the dynamics of the membrane and the protein. For the wash-out experiments, the well was washed 5 times with 30 µl wash buffer before another FRAP experiment was performed.

#### Peptide experiments on SLBs

After carefully washing the access protein, the peptides were incubated for 30 min to allow for binding to mCherry-hIRE1α LD-10His. Phase separation was induced with 25 mM Hepes pH 7.3, 150 mM NaCl, 250 µM TCEP and PEG at percentages between 9 % and 11 %. After an 10 min incubation period, the Atto488 labeld membrane and the mCherry tagged protein were imaged. The end concentration of the peptides above the SLB were 10 µM MPZ1N, 1 µM MPZ1N −2X and 1 µM MPZ1N −2X-RD.

#### In solution phase separation assay

The phase separation behavior of hIRE1α LD protein variants in presence of PEG (Sigma-Aldrich (P2139) or 40 % (w/w) Sigma-Aldrich (P1458)) was observed *via* DIC microscopy on a Zeiss Axio Observer inverted microscope. Images were acquired at room temperature with a Plan-Apochromat 63x/1.4 Oil DIC RMS objective and CoolSnapHQ2 or Hamamatsu ORCA-Flash4.0 LT+ Digital CMOS camera controlled by Visitron and Zeiss systems, respectively. Therefore, glass wells (Greiner Bio-One 96 Well SensoPlate™) were pretreated with 1 % (w/v) Pluronic® F-127 (PF127, Sigma Aldrich) for 2 hrs at room temperature. After three wash steps (150 mM NaCl, 1 M Hepes pH 7.3 (Molecular Biology, Fisher BioReagents™), phase separation of the protein of interest was induced. Hence, the protein was mixed with equal volumes of PEG containing buffer (25 mM HEPES pH 7.3, 150 mM NaCl, 4 mM DTT, 20 mM MgCl2, 2X PEG percentage (depending on condition)) in a final volume of 50 µl. For peptide experiments, 24.5 µM hIRE1α LD and 0.5 µM mCherry-hIRE1α LD-10His were preincubated with the respective peptide for 30 min on ice before phase separation was induced *via* PEG. The final protein and PEG concentration and incubation time is indicated in the Figure legends. Image processing was performed in ImageJ and Adobe Photshop® adjusting the brightness, contrast and sharpness of the images.

### FRAP on condensates in solution

Phase separation was induced as described in the “In solution phase separation” section. For FRAP experiments hIRE1α LD was mixed with 2 % of the corresponding mCherry tagged protein at a concentration of 25 µM, phase separation was induced in a test tube for 30 min in the presence of 6 % PEG in the well. Experiments were performed on a Zeiss Axio Observer inverted microscope equipped with a Yokogawa CSU-X1-A1 Nipkow spinning disc unit (Visitron Systems; pinhole diameter 50 μm, spacing 253 μm), sCMOS camera (Pco.edge 4.2) and a Plan-Apochromat 63x/1.4 Oil DIC objective. Images were conducted every 5 s for a time course of 10 min with 80 % HX, 50 ms exposure and 10 % 561 nm laser intensity exposed for 100 ms. Per condition, 3 condensates were bleached after 2 frames with 100 % 561 nm laser power for 10 ms per pixel. Image processing was performed in ImageJ selecting the FRAP ROI and two ROIs of the same size within a non FRAPed condensate for bleaching correction and an area without condensate for background correction for every FRAPed condensate. The background value was subtracted from the FRAP and bleaching value, followed by calculating the bleaching factor to correct the FRAP values leading to the final normalization. The normalized FRAP values were fitted to the One-phase association in PRISM.

#### Recruitment experiments

Phase separation was induced as described in the “In solution phase separation” section. After 30 min of incubation within the well, the respective mCherry labeled protein was added (at 2 % of a total protein concentration of 25 µM) and imaged under the same conditions (on a Zeiss Axio Observer inverted microscope equipped with a Yokogawa CSU-X1-A1 Nipkow spinning disc unit (Visitron Systems; pinhole diameter 50 μm, spacing 253 μm), sCMOS camera (Pco.edge 4.2) and a Plan-Apochromat 63x/1.4 Oil DIC objective) every 5 s for 25 min with 80 % HX, 50 ms and 10 % 561 laser intensity exposed for 100 ms. Image processing was performed in ImageJ.

### Modelling of disordered regions

The protein structure of the human IRE1α core Lumenal Domain (cLD) dimer was obtained from the Protein Data Bank (www.rcsb.org ^64^, PDB ID: 2HZ6) ^21^. We added the missing residues (66-70, 89-90, 11-115, 131-152, 308-357, 369-443) as unfolded loops using UCSF Chimera (version 1.15, ^65^

From this model, the regions DR1, DR2 and linker were extracted as isolated peptides and individually mapped to coarse-grained representation.

The peptides obtained were:

**DR1,** 131 – LTGEKQQTLSSAFADSLSPSTS - 152;

**DR2**, aa 307-VPRGSTLPLLEGPQTDGVTIGDKGESVITPSTDVKFDPGLKSKNKLNYLRNY-358;

**Linker region**, aa 369 - LSASTKMLERFPNNLPKHRENVIPADSEKKSFEEVINLVDQTSENAPTTVSRDVEEKPAHAPARPEAPVDSMLKD - 443.

The conversion of the all-atom models into Martini 3 ^66^ coarse-grained models and the setup of the simulation systems were performed using the tools *martinize2* (https://github.com/marrink-lab/vermouth-martinize) and *insane.*py ^67^ and gromacs/2020.5 tools (*gmx insert-molecules*). The termini were neutralized and the side chain fix was applied to prevent unrealistic side chain orientations as proposed in ^68^.

### Molecular dynamics simulations

We set up systems containing two disordered regions’ peptides by randomly inserting 16 copies of each region in a 30 x 30 x 30 nm^3^ simulation box. We obtained a system containing DR1 and DR2 and a system containing DR2 and the linker region. We solvated the systems with Martini water molecules and chloride and sodium ions, corresponding to a salt concentration of 150 mM.

After a first energy minimization we equilibrated the system. First, we ran a 10 ps-long simulation using a 1 fs time step and restraining the position of protein backbone beads by using harmonic potentials with force-constants of 1000 kJ mol^−1^ nm^−2^. Afterwards, we ran another 2.1 ns without restraints using a 30 fs time step and a final equilibration of 21 ns. After the equilibration, we ran MD simulations using 20 fs time step. The temperature in the simulation box was controlled by a velocity rescale thermostat ^69^(reference temperature T_ref = 300K, coupling time constant tau_T = 1 ps). The Parrinello-Rahman barostat ^70^ (reference pressure p_ref = 1 bar; coupling time constant τ_p = 24 ps) was used for the last equilibration step and for the production run.

Coarse-grained molecular dynamics simulations were performed using with the Martini 3.0 forcefield ^66^ and the GROMACS 2020.5 software ^71^.

### Contact maps from MD simulations

We set up individual simulations for each region (DR1, DR2 and linker) in two different settings, namely containing two peptides or 33 peptides. For the two peptides’ simulations, we determined the dimensions of the box by setting a 2 nm distance between periodic images in a cubic box. In the latter simulation setting we randomly inserted 33 peptide copies in a 30 x 30 x 30 nm^3^ simulation box to obtain a protein concentration of 2 mM. We solvated the systems with Martini water molecules and chloride and sodium ions, corresponding to a salt concentration of 150 mM.

We analyzed the contacts formed over time among the peptide chains in these simulations. Initially, we computed the contact map between all beads of all peptides at each frame thanks to the python package Contact Map Explorer (https://github.com/dwhswenson/contact_map, version 0.7.0). Two Martini beads were considered in contact if nearer than 0.5 nm. In the simulations containing 33 copies, we considered which peptide chains are interacting to create a network representation of the clusters at each frame from which we could determine which is the central chain of the cluster. Then we counted all the contacts between beads of the central chain and beads of its neighboring chains at each frame and we averaged over the number of chains interacting with the central one at each frame.

We obtained a matrix of dimensions (*Number of beads per chain*, *Number of beads per chain*) and we convert it to dimensions (*Number of residues per chain*, *Number of residues per chain*) by retaining the maximum score present between all the beads of a pair of residues. The contact matrices were computed in a similar way for systems of two peptides. In these simulations we considered all the interactions happening between the two chains, removing the notion of a central chain. We produced 1D-projections of the contact maps by summing up all the contribution for a specific residue in the final contact matrices for simulations of two or 33 copies.

### Generation of Constructs for Stable Cell Lines *via* Lentiviral Transduction

For the establishment of stable cell lines in Mouse Embryonic Fibroblasts, a vector with a Tet-On doxycycline-inducible TRE3G promoter was utilized. TRE3G-P2A-eBFP2-PGK-puroSTOP-IRES-rtTA3 (kind gift from Gijs Versteeg) was cut using restriction enzymes BsrGI-HF and BamHI (New England Biolabs). hIRE1α signal sequence with the transmembrane domain (amino acids 1-469) 3XFlag and 6XHis tag and hIRE1α kinase-RNase domain (amino acids 470-977) were amplified frompShuttle-CMV-TO_hsIRE1-3F6H-GFP-LKR-K36.3. Additionally, due to its higher stability, the GFP tag, upstream of the kinase domain, was replaced with mNeonGreen through Gibson Assembly. The mutations were introduced into IRE1α N-terminal part (amino acids 1-469) or into kinase-RNase domain (amino acids 470-977) *via* PCR and the mutated fragment were used for Gibson assembly as described above. All the constructs, except for the core LD, encode for the full-length lumenal sequence of hIRE1α (amino acids 1-469). The lumenal boundaries for the cLD include the core sequence (amino acids 24-389), as well as, a short region proximal to the transmembrane domain (amino acids 434-443), which was shown to be essential for the interaction with the Sec61 translocon ^72^.

### Transfection of Packaging Cells

All transfections were performed by mixing DNA and Polyethylenimine (PEI, Polysciences, 23966) in a 1:3 ratio (μg DNA/μg PEI) in DMEM without supplements. Plasmids for the transfection were purified using an endotoxin-free Plasmid Kit (Qiagen). Transfection was performed using 1100 ng of total DNA (500 ng transfer plasmid, 500 ng pCMVR8.74 Addgene plasmid # 22036, 100 ng pCMV-VSV-G Addgene plasmid # 8454) The day before transfection, 2*10^5^ HEK293T HiEx packaging cells were seeded in 6-well plates in fully supplemented media. The following day, the above-described transfection mixture was added dropwise to the cells. Subsequently, cells were incubated for 48 hrs.

### Transduction of Mouse Embryonic Fibroblasts (MEF) (IRE1α^−/−^/IRE1β^−/−^) and Cell Selection

Following the 48 hour incubation period, the viral supernatant was sterile filtered with a syringe. The day before transduction 1*10^5^ MEF (IRE1α^−/−^/IRE1β^−/−^) were seeded in a 6-well plates in fully supplemented media. For the transduction, the virus was mixed with fully supplemented DMEM (Sigma-Aldrich, D6429) and 8 μg/ml Polybrene (Sigma-Aldrich, TR-1003-G) at 1:50 (v/v). After a 48 hour incubation period, cell lines were expanded to 10-15 cm dishes. Protein expression, for subsequent Fluorescence Activated Cell Sorting (FACS), was induced with 400 nM of Doxycycline for 24 hours. Cells were sorted in yield mode using BD FACSAria II or BD FACSMelody, gated for low and high-expression cells. The high-expression cells were resorted in stringent mode, following the same procedure. The second FACS sorted high population of the first FACS sorted high population was used for characterization.

### Immunofluorescence

IRE1 double-knockout Mouse Embryonic Fibroblasts (MEF) (IRE1 α^−/−^/IRE1β^−/−^) reconstituted with a doxycycline inducible hIRE1α-mNG (or mutants) were seeded at a density of 20000 cells in a µ-Slide 8well dish (ibidi) 1 day before the experiment. IRE1α expression was induced for 24 hrs by adding 400 nM doxycycline. Cells were stressed with 5 µg/ml Tunicamycin for 4 hrs. The experiment was stopped by washing with cold PBS and fixation with 4 % paraformaldehyde for 7 min. After two more washes with PBS, the cells were incubated for 1h in blocking buffer (PBS, 10 % FBS, 1 % Saponin) followed by primary antibody incubation overnight at 4°C (Calnexin, Abcam ab22595 at a dilution of 1:200). After washing twice with wash buffer (PBS, 10 % FBS) the secondary antibody (Alexa Fluor 594 goat anti-rabbit, Invitrogen A11037 at a dilution of 1:1000) was incubated for 1h at room temperature. After three additional wash steps, the sample was imaged in PBS on a Zeiss LSM 980 inverse point scanning confocal microscope with a Plan-Apochromat 63x/1.4 Oil DIC, WD 0.19 mm objective. The microscope is operated by the Zeiss ZEN 3.3 microscope software. mNG and Atto549 were excited by the 488 nm and 561 nm laser diodes of the microscope, respectively. Image processing was performed in ImageJ.

### Western blotting

Treated MEFs at a confluency of 80 % were collected in RIPA buffer. The protein concentration was determined by a bicinchoninic acid assay using a commercially available kit. 10-15 μg protein of the lysate in sample buffer was loaded after denaturation for 10 min at 95°C on a 10 % sodium dodecyl sulfate gel. The proteins were wet transferred from the gel to a nitrocellulose membrane in transfer buffer (25 mM Tris, 192 mM glycine, 20 % (v/v) ethanol, pH 8.3) for 120 min at 110 V. The proteins on the membrane were stained with Ponceau S for 5 min followed by blocking in 5 % milk for 1 h at room temperature. The primary antibody was applied in 2.5 % milk for 1 h at room temperature or overnight at 4°C. The membranes were washed five times in TBST for 5min before the secondary antibody in 2.5 % BSA (Anti-Rabbit IgG (H+L), HRP Conjugate, Promega W401B at a dilution of 1:10000) was added and incubated for 1h at room temperature. After five 5 min TBST wash steps, the chemiluminescence substrate for the horseradish peroxidase was applied using a commercially available kit. The membranes were imaged using a ChemiDoc system and analyzed with the Image Lab software of Bio-Rad.

Primary antibodies

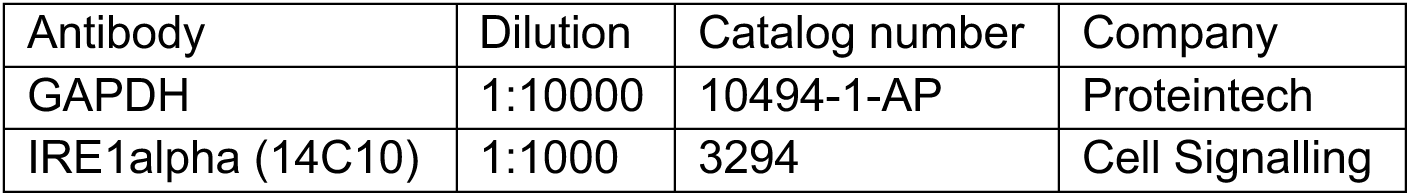

### *XBP1* mRNA splicing assays

#### Semi quantitative PCR analyses

The protocol was adapted from Karagöz et al 2017, elife.^22^ In brief, MEFs grown in a 12 well plate were treated for 24 hrs with or without 400 nM dox, DMSO or tunicamycin (5 µg/ml) and collected in 180 µl TriFast (VWR Life Science). 100 µl of water and 60 µl of chloroform was added, mixed and incubated for 10 min at room temperature followed by a 5 min 20,800 x g spin. The transparent phase was transferred to a new tube, mixed with 100ul isopropanol and 0.5ul glycogen and incubated for 15 min on ice. After a 10 min 20,800 x g spin, the pellet was washed three times with 75 % ethanol and resuspended in 16 µl water. The total RNA concentration was determined by Nanodrop measurement and normalized throughout the samples. The quality of RNA was verified by a 1 % Agarose gel. To generate cDNA, total RNA (a minimum of 175 ng) was reverse transcribed using LunaScript RT (New England Biolabs) followed by dilution of 1:5 or 1:10 depending on the normalized RNA input concentration. 4 % cDNA product was used to perform semiquantitative PCR using 50 % Taq MM (New England Biolabs) and 0.5 µM of the forward (GAACCAGGAGTTAAGAACACG) and reverse (AGGCAACAGTGTCAGAGTCC) primers. The PCR product was amplified for 28 cycles and analyzed on a GelRad stained 3 % agarose gel (50:50 mixture of regular and low-melting point agarose). The gels were imaged using a FastGene FAS_V Geldoc System and analyzed with the Image Lab software of Bio-Rad.

#### Real-time quantitative reverse transcription PCR analyses

The qPCRs were conducted on a Roche LightCycler^®^ 480 in 384 well plates in triplicate. Per well there was 2ul cDNA, 0.8ul of Forward and Reverse primer (10uM concentration) each, 5ul Promega GoTaq® qPCR mix (cat.# A6002) and 2.2ul nuclease/RNase free water to a total volume of 10ul. Data was processed using the ΔCq method in R with the tidyqpcr package ^73^. The values are plotted as relative fold change of the target normalized to their respective reference gene expression level. Target primers for spliced Xbp1 were 5′-CTGAGTCCGAATCAGGTGCAG-3′ for forward and 5′ GTCCATGGGAAGATGTTCTGG-3′ for reverse, taken from Scortegagna et al. ^74^. The reference gene for normalization was mHPRT (Hypoxanthine guanine Phosphoribosyl-transferase) (Forward primer: 5′ GCAGTCCCAGCGTCGTGATTA-3′, Reverse primer: 5′-TGATGGCCTCCCATCTCCTTCA-3′) from Manakanatas et al. ^75^.

## Figures

**Fig. Supp. 1.**
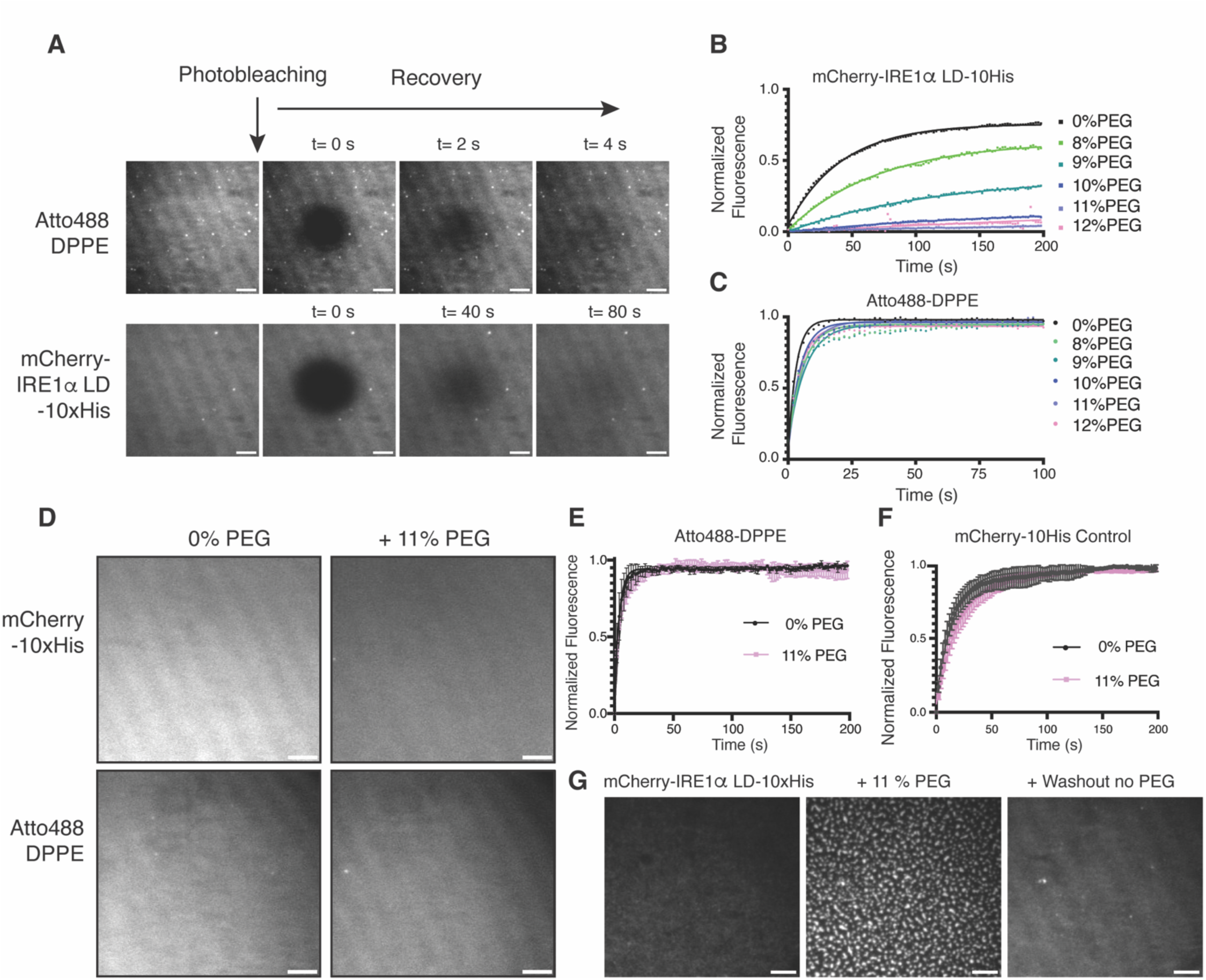
**A.** TIRF images of FRAP experiments of Atto488 labeled DPPE lipids (top) and mCherry-IRE1α LD-10His (bottom) on SLBs showing the dynamic behavior within the indicated time. Scale bar = 5 µm. **B.** FRAP curves of mCherry-IRE1α LD-10His tethered to SLBs by 1 % Ni-NTA labeled lipids. The SLBs are incubated 10 min with the indicated concentration of the crowding agent PEG before the images are taken. The mobile fraction and diffusion values are decreasing with increasing PEG concentration. **C.** FRAP curves displaying the fluorescent intensity of Atto488 labeled DPPE lipids within SLBs treated with the indicated concentration of the crowding agent PEG over time. **D.** TIRF images displaying mCherry-10His control and the membrane (Atto488 DPPE) with and without PEG. Scale bar = 5 µm. **E.** FRAP curves displaying the fluorescent intensity of Atto488 labeled DPPE lipids within SLBs treated with the indicated concentration of the crowding agent PEG over time. **F.** FRAP curves displaying the fluorescent intensity of mCherry-10His control within SLBs treated with the indicated concentration of the crowding agent PEG over time. **G.** TIRF images of mCherry-hIRE1α LD-10His tethered to SLBs by 1 % Ni-NTA labeled lipids in the absence of PEG, in presence of 11 % PEG and where PEG is washed out from the well. Scale bar = 5 µm.

**Fig. Supp. 2.**
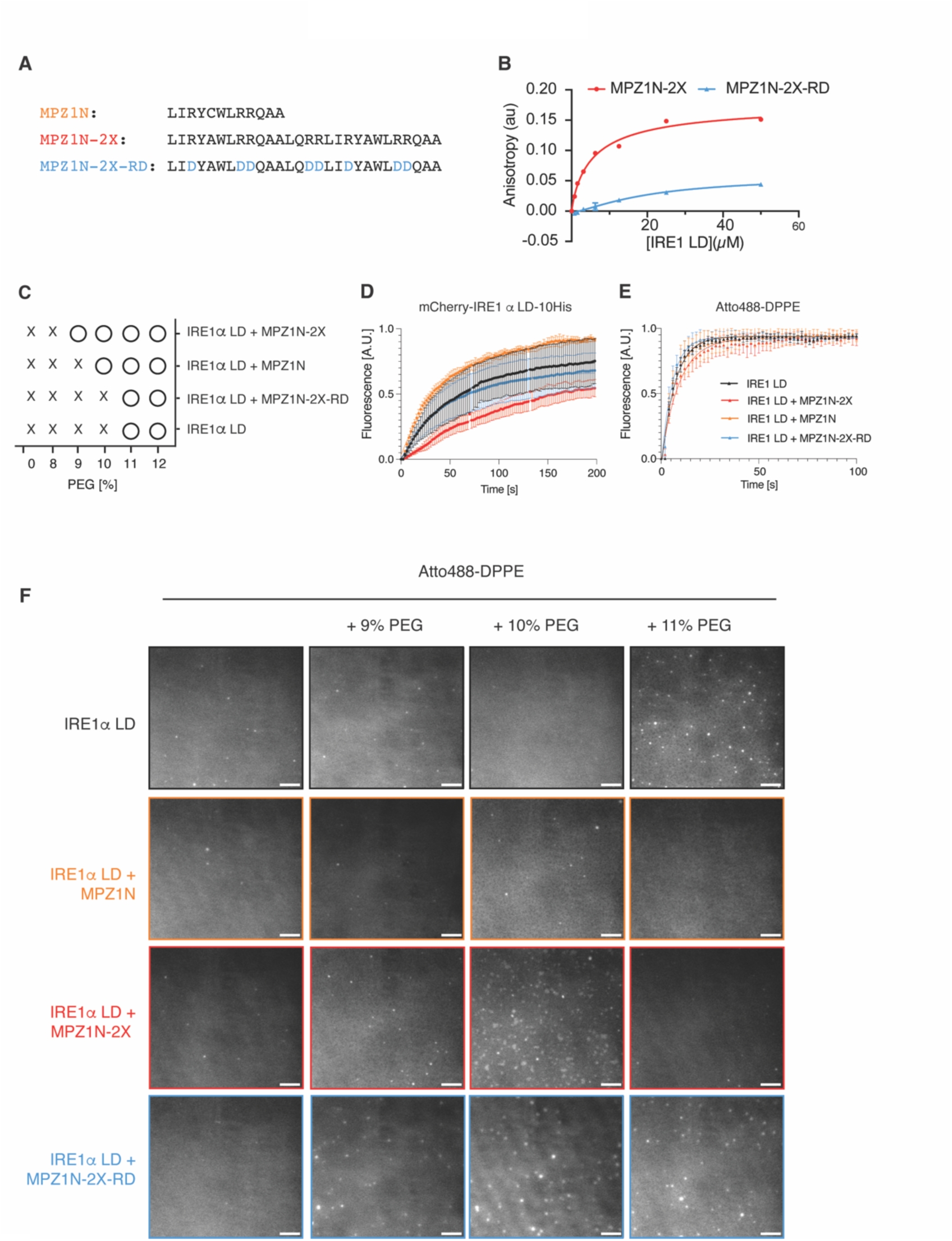
**A**. Amino acid sequences of model unfolded polypeptides MPZ1N and MPZ1-N-2X and the control non-binding derivate MPZ1N-N-2X-RD. **B.** Fluorescence anisotropy experiments monitor the interaction of N-terminal fluorescein labeled MPZ1N-2X and its derivative MPZ1N-2X-RD with IRE1α LD. MPZ1N-2X interacts with IRE1α LD at 2 µM affinity, whereas the MPZ1N-2X-RD is impaired in binding. **C.** Diagram summarizing mCherry-IRE1α LD-10His clustering on SLBs in the presence of peptides at various PEG concentrations. “X” depicts no cluster and “O” cluster formation. **D.** FRAP curves of mCherry-IRE1α LD-10His on SLBs in the absence (black curve) and presence of 10 µM MPZ1N (orange curve), 1 µM MPZ1N-2X (red curve) and 1 µM MPZ1N-2X-RD (blue curve) peptides. Curve marks show the mean value, error bars display the standard deviation and the values are fitted to a one-phase association curve. n=3 independent experiments were performed. **D**. FRAP curves of Atto488 labeled DPPE lipids within SLBs color code is as in **Fig. Supp. 2D**. **E.** TIRF images displaying Atto488 labeled DPPE within SLBs of experiments shown in **Fig. 1F-I**. Scale bar = 5 µm

**Fig. Supp. 3.**
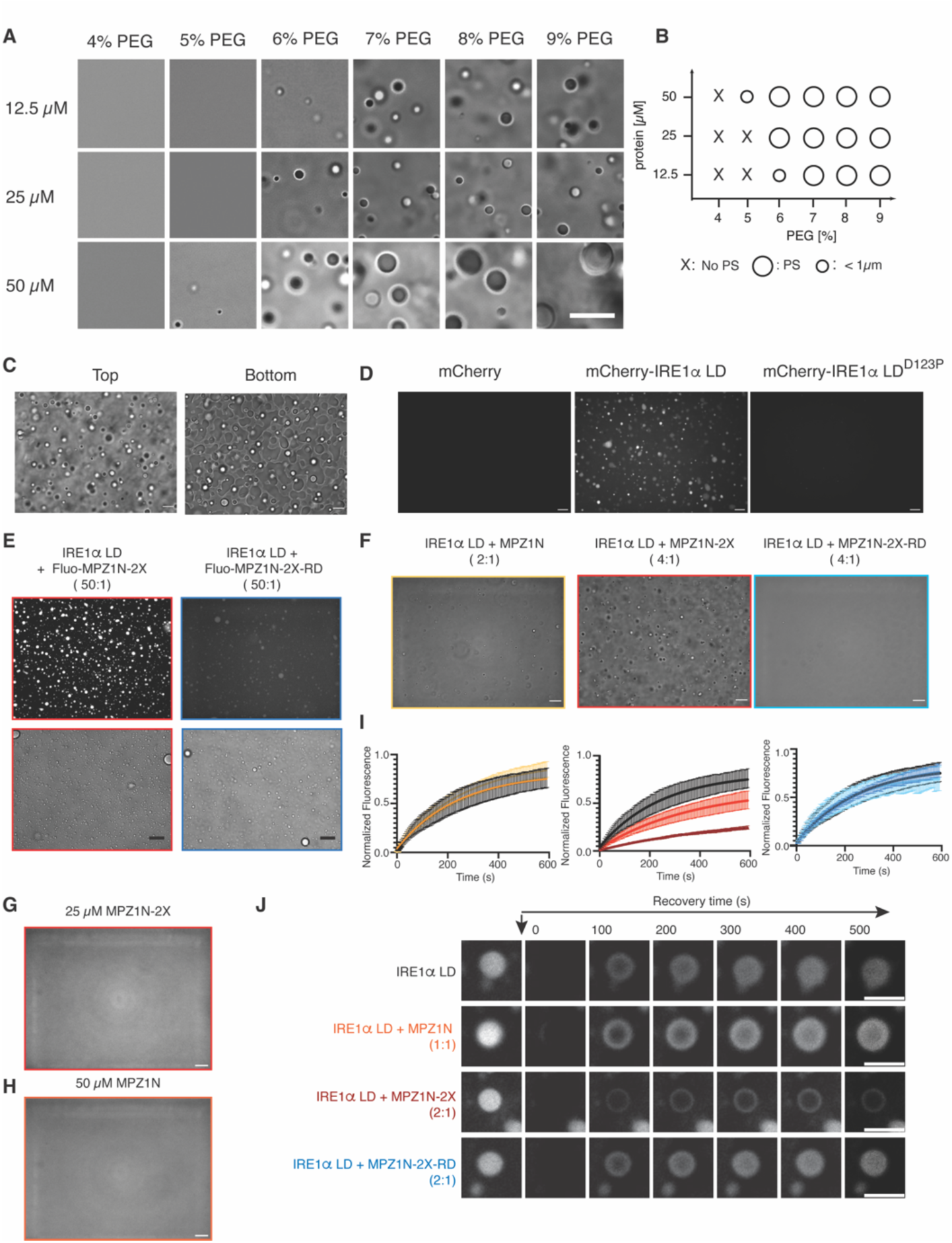
**A.** DIC images of IRE1α LD representing the phase diagram of IRE1α LD. Scale bar = 10 µm. **B.** Phase diagram of IRE1α LD condensates at 12.5, 25 and 50 µM at 30 min incubation with 4-9 % PEG as in **Fig. Supp. 1A**. No phase separation (PS) is indicated by a cross and phase separation (PS) is indicated by a circle. The smaller circle refers to condensates with diameter < 1 µm. **C.** DIC images of 50 µM IRE1α LD incubated with 6 % PEG for 30 min. The images are obtained at the bottom or middle of the plate. Scale bar = 10 μm. **D.** Fluorescence images of 25 µM mCherry-10His control, mCherry-IRE1α LD-10His and the dimerization mutant of IRE1α LD, mCherry-IRE1α LD^D123P^-10His after 30 min incubation with 6% PEG. Scale bar = 10µm. **E.** Confocal (top) and bright field (bottom) images displaying the recruitment of Fluorescein-labeled MPZ1N-2X (left, red) and MPZ1N-2X-RD (right, blue) peptides into preformed IRE1α LD condensates. Scale bar = 13 μm. **F.** DIC microscopy images of 50 µM IRE1α LD incubated with MPZ1N (2:1 stoichiometry, left), MPZ1N-2X (4:1 stoichiometry, middle) or MPZ1N-2X-RD (4:1 stoichiometry, right panel) at 30 min after induction of phase separation with 5% PEG. Scale bar = 10 μm. **F.** DIC images of 25 µM MPZ1N-2X peptide in the presence of 6 % PEG. Scale bar = 10 μm **G**. DIC images of 50 µM MPZ1N peptide in the presence of 6 % PEG. Scale bar = 10 μm **H.** FRAP curves of 25 µM IRE1α LD and 6 % PEG in the absence (black curve) and in the presence of MPZ1N peptide (2:1 stoichiometry, light orange curve, 1:1 stoichiometry orange curve), MPZ1N-2X peptide (4:1 stoichiometry, red, 2:1 stoichiometry dark red) and MPZ1N-2X-RD control peptide (4:1 stoichiometry, light blue, 2:1 stoichiometry blue). **I.** FRAP images of a single IRE1α LD condensate in absence and presence of the model unfolded peptides at the indicated stichometry taken before and at the indicated time points after photo-bleaching. Scale bar = 5 µm.

**Fig. Supp. 4.**
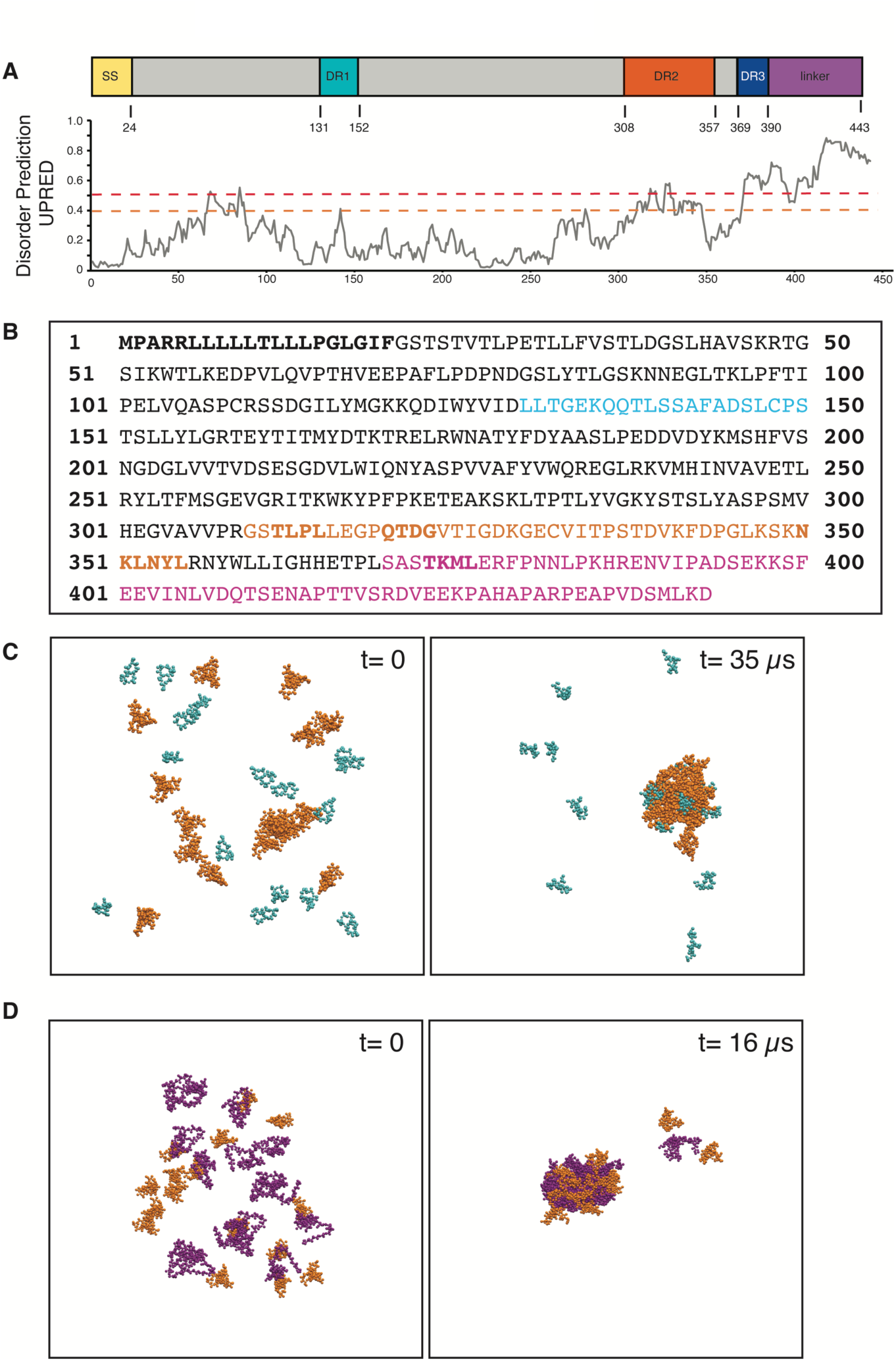
**A.** Schematic presentation of DRs in IRE1α LD domain organization in combination with the prediction of intrinsically unstructured regions of hIRE1α LD using the IUPRED server. The red and orange lines indicate moderate and disordered propensity, respectively. **B.** Amino acid sequence of IRE1α LD where DR1, DR2 and linker segments are colored in cyan, orange and purple respectively. The signal sequence and mutated segments are highlighted in bold letters. **C**. Simulation of 16 copies of DR1 (cyan) and 16 copies of DR2 (orange). Molecular Dynamics Simulations show that the cluster formed by DR2 do not recruit DR1 segments. **D.** Simulation of 16 copies of DR2 (orange) and 16 copies of linker region (purple) (right two panels). Molecular Dynamics Simulations show that the cluster formed by DR2 recruit the linker segments.

**Fig. Supp. 5.**
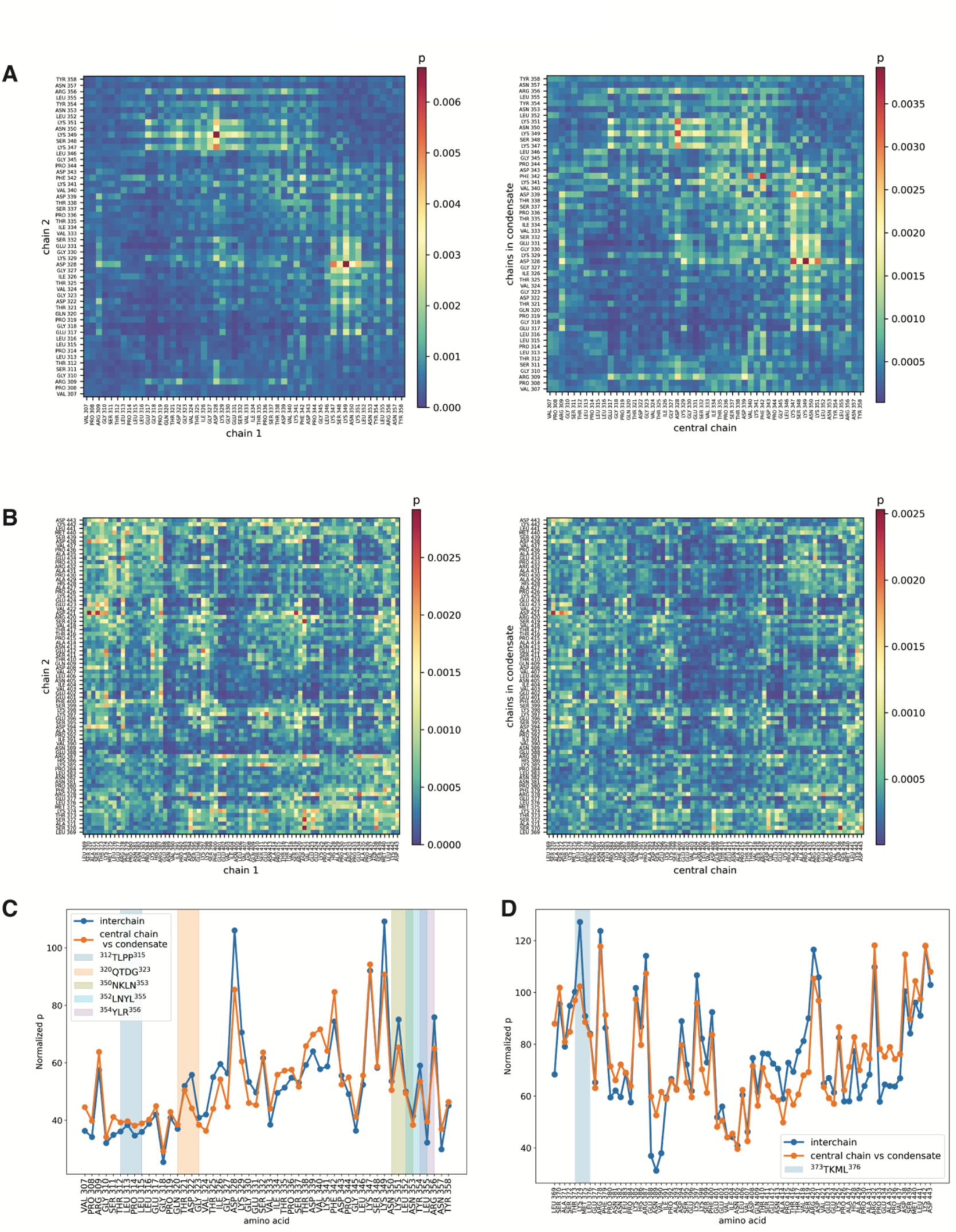
**A.** Contact maps for interchain and condensate interactions in simulations containing two or 33 copies of DR2. **B.** Contact maps for interchain and condensate interactions in simulations containing two or 33 copies of linker. **C.** 1D-projections of the contact maps computed for the simulations containing two or 33 copies of DR2. **D.**1D-projections of the contact maps computed for the simulations containing two or 33 copies of linker.

**Fig. Supp. 6.**
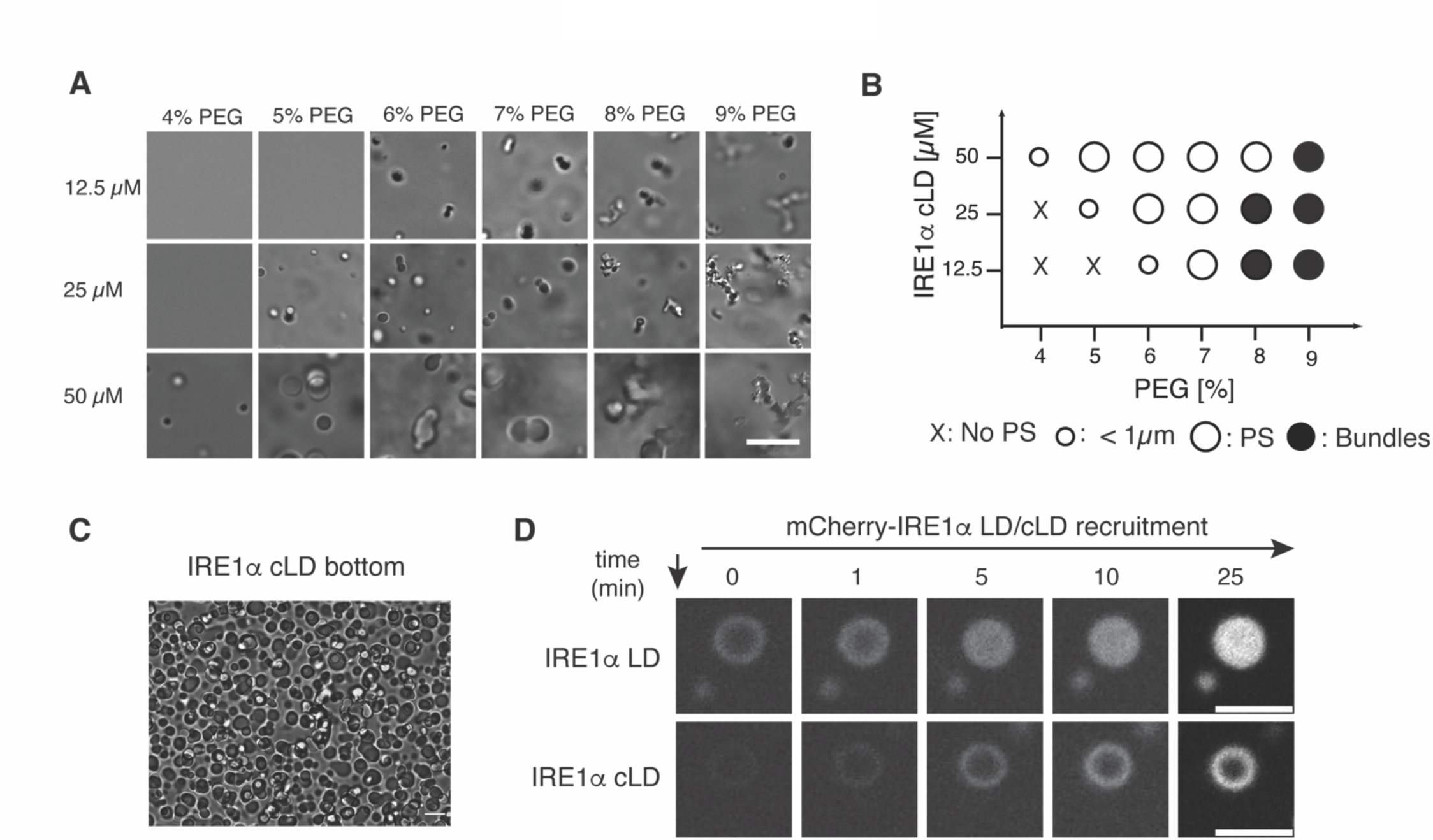
**A.** DIC Images of IRE1α cLD representing the phase diagram at 12.5, 25 and 50 µM acquired after 30 min incubation with PEG at concentrations ranging from 4 - 9 %. Scale bar = 10 µm **B.** Phase diagram of IRE1α cLD based on images in Fig. Supp. 4C. No phase separation (PS) is indicated by a cross, phase separation (PS) is indicated by a circle and condensates that resemble beads on a string are represented by a black circle (bundles). The smaller circle refers to smaller condensates (diameter < 1 µm). **C.** DIC images of the bottom of the well of IRE1α cLD (50 µM) condensates taken 60 min after induction of phase separation *via* addition of 6 % PEG showing the phase separation propensity and wetting effect. Scale bar = 10 μm **D.** Fluorescence images of 25 µM IRE1α LD (top) or IRE1α cLD (bottom) condensates at the indicated time points after 30 min incubation with 6 % PEG following the recruitment of 2 % mCherry labeled IRE1α LD or cLD, respectively. mCherry-IRE1α LD-10His is recruited to the center of preformed IRE1α LD condensates, whereas mCherry-IRE1α cLD-10His could only associate with the outer shell of the preformed IRE1α cLD condensates. Scale bar = 5 µm.

**Fig. Supp. 7.**
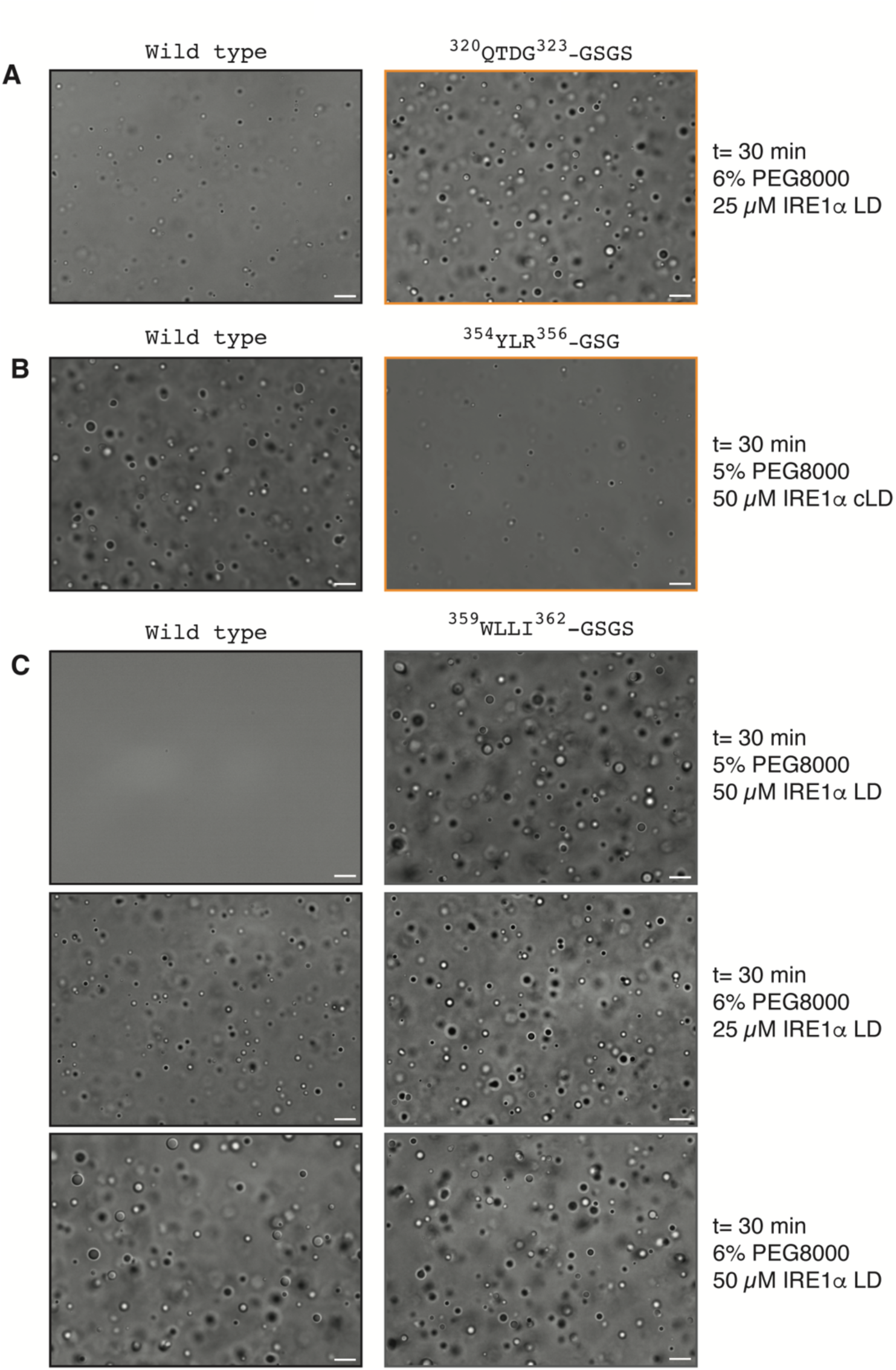
**A.** DIC images of 25 µM WT IRE1α LD and IRE1α LD ^320^QTDG^323^-GSGS mutant showing LLPS behavior 30 min after induction of phase separation by the addition of 6 % PEG. **B.** DIC images of 50 µM WT IRE1α cLD and IRE1α cLD ^354^YLR^356^-GSG mutant showing LLPS behavior 30 min after induction of phase separation by the addition of 5 % PEG. **C.** DIC images comparing the LLPS behavior of WT IRE1α LD (left column) and IRE1α LD ^359^WLLI^323^-GSGS mutant (right column) 30 min after induction of phase separation at 50 µM protein concentration and 5 % PEG (top row) at 25 µM protein concentration and 6 % PEG (middle row) and at 50 µM protein concentration and 6 % PEG (bottom row). Scale bar for all images = 10 μm.

**Fig. Supp. 8.**
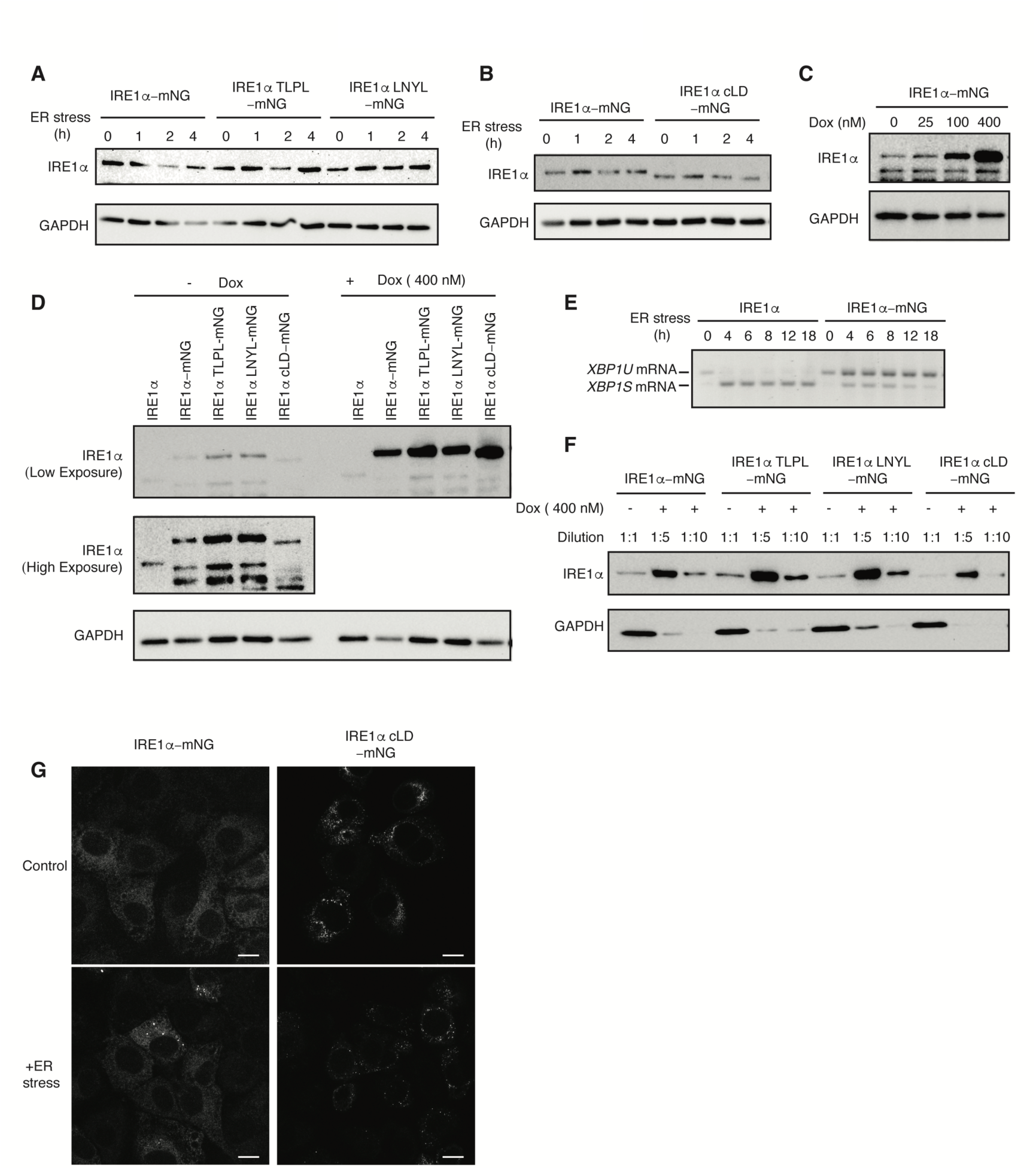
**A,** Western blot analyses comparing the expression level of IRE1α-mNG and its mutants IRE1α TLPL-mNG and IRE1α LNYL-mNG in MEFs in the absence of doxycycline treatment at different points after induction of ER stress. **B.** Western blot analyses comparing the expression levels of IRE1α-mNG and IRE1α cLD-mNG in MEFs in the absence of doxycycline treatment at different points after induction of ER stress. **C.** Western blot analyses comparing the WT IRE1α expression to the expression level of IRE1α-mNG in the absence and presence of various concentrations of doxycycline inducing its expression for 24 hours. **D.** Quantification of overexpression levels of IRE1α-mNG and its mutants upon induction of protein expression with 400 nM doxycycline for 24 hrs. **E.** Semiquantitative PCR reaction to monitor splicing of *XBP1* mRNA by IRE1α-mNG (in the absence of doxycycline) and wild type IRE1α at different time points after induction of ER stress by addition of 5 µg/ml Tunicamycin. The bands are indicated as unspliced and spiced *XBP1* variants. **F.** Western blot analyses comparing the expression levels of IRE1α-mNG and its mutants in MEFs in the absence and presence of 400 nM doxycycline treatment. Lysates obtained from 400 nM doxycycline MEFs were diluted 1 to 5 and 1 to 10. **G.** Immunofluorescence images of MEFs treated with 100 nM doxycycline to induce expression of IRE1α-mNG and the IRE1α cLD-mNG mutant in the absence (top row) of stress and treated with 5 µg/ml ER stressor Tunicamycin for 4 hrs (bottom row). IRE1α-mNG and its mutants are visualized by mNG fluorescence (green). Scale bar = 10 µm.

