## Supplemental Tables for "Stress-induced clustering of the UPR sensor IRE1α is driven by disordered regions within its ER lumenal domain"

Kettel et al., **Supp.Table 1 .**

**mCherry-IRE1 LD-10His** Fluorescence Intensity on the SLBs

| Concentration of Ni-NTA Lipid ( mol %) | Concentration of mCherry-IRE1 LD-10His | Fluorescence Intensity (AU) |
|---|---|---|
| 1% | 50 nM | 2655.90 |
| 1% | 200 nM | 9267.75 |
| 1% | 500 nM | 12421.96 |
| 1% | 1 μM | 13245.89 |
| 2% | 50 nM | 4994.50 |
| 2% | 200 nM | 13733.22 |
| 2% | 500 nM | 18802.05 |
| 5% | 1 μM | 27346.64 |

Kettel et al., **Supp. Table 2.** linked to the data in **Fig. 1** and **Fig. Supp. 1 and Fig. Supp 2.**

Fits of the FRAP curves of **mCherry-hIRE1 LD-10His** on SLBs in **Fig 1F-I**

| mCH | **IRE1 LD** | **+ MPZ1N** | **+MPZ1N-2X** | **+ MPZ1N-2X-RD** |
|---|---|---|---|---|
| **Fit values y = a*(1-exp(-b*x))** | | | | |
| Mobile fraction [%] | 76.48 ± 14.45 | 89.77 ± 3.07 | 66.80 ± 4.52 | 81.93 ± 14.32 |
| Half-life time [s] | 37.64 ± 6.64 | 27.60 ± 4.51 | 83.25 ± 24.00 | 32.95 ± 6.81 |
| Diffusion coefficient [μm$^2$/s] | 0.08 ± 0.04 | 0.12 ± 0.02 | 0.03 ± 0.01 | 0.09 ± 0.04 |

Fits for the FRAP curves of **Atto488-DPPE** lipids within SLBs in **Fig. Supp. 2E** where **mCherry-hIRE1 LD-10His** is attached in **Fig 1F-I**

| | Atto488-DPPE | +MPZ1N | +MPZ1N-2X | +MPZ1N-2X-RD |
|---|---|---|---|---|
| **Fit values y = a*(1-exp(-b*x))** | | | | |
| Mobile fraction [%] | 97.25 ± 2.28 | 93.53 ± 3.17 | 96.30 ± 1.96 | 94.18 ± 2.87 |
| Half-life time [s] | 2.78 ± 0.5 | 2.36 ± 0.23 | 2.95 ± 0.88 | 2.35 ± 0.52 |
| Diffusion coefficient [μm$^2$/s] | 1.04 ± 0.41 | 1.22 ± 0.47 | 0.95 ± 0.14 | 1.21 ± 0.39 |

Fits for the FRAP curves of **mCherry-hIRE1 LD-10His** on SLBs in the absence and presence of various PEG concentrations**. Fig. Supp. 1B**

| | 0%PEG | 8%PEG | 9%PEG | 10%PEG | 11%PEG | 12%PEG |
|---|---|---|---|---|---|---|
| **Fit values y = a*(1-exp(-b*x))** | | | | | | |
| Mobile fraction [%] | 69.70 | 57.00 | 46.00 | 23.2 | 11.9 | nan |
| Half-life time [s] | 25.20 | 47.31 | 87.67 | 126.80 | 291.39 | nan |
| Diffusion coefficient [μm$^2$/s] | 0.18 | 0.10 | 0.05 | 0.04 | 0.02 | nan |

Fits for the FRAP curves of **Atto488-DPPE** within SLBs in the absence and presence of various PEG concentrations**. Fig. Supp. 1C**

| | 0%PEG | 8%PEG | 9%PEG | 10%PEG | 11%PEG | 12%PEG |
|---|---|---|---|---|---|---|
| **Fit values y = a*(1-exp(-b*x))** | | | | | | |
| Mobile fraction [%] | 97.4 | 96.8 | 96.6 | 96.4 | 96.3 | 96.2 |
| Half-life time [s] | 2.20 | 2.57 | 2.76 | 2.65 | 3.00 | 2.63 |
| Diffusion coefficient [μm$^2$/s] | 2.09 | 1.78 | 1.66 | 1.73 | 1.53 | 1.74 |

Fits for the FRAP curves of **Atto488-DPPE** lipids and **mCherry-10His** control on SLBs. **Fig. Supp. 1E,F**

| | Membrane **Atto488-DPPE** | | Control Protein **mCherry-10XHis** | |
|---|---|---|---|---|
| | - | **+ 11%PEG** | - | **+ 11%PEG** |
| **Fit values y = a*(1-exp(-b*x))** | | | | |
| Mobile fraction [%] | 93.07 ± 3.84 | 98.07 ± 1.11 | 94.57 ± 2.91 | 95.53 ± 1.40 |
| Half-life time [s] | 1.26 ± 0.07 | 1.77 ± 0.32 | 10.38 ± 2.88 | 15.47 ± 4.93 |
| Diffusion coefficient [μm$^2$/s] | 3.63 ± 0.20 | 2.65 ± 0.50 | 0.47 ± 0.14 | 0.32 ± 0.09 |

Kettel et al., **Supp. Table 3.** linked to the data in **Fig. 2C,D, Supp. Fig 3I,J**

Fits of the FRAP curves of **mCherry-IRE1 LD-10His** in condensates formed in solution

| | hIRE1 LD | hIRE1 LD + MPZ1N (1:1) | hIRE1 LD + MPZ1N (2:1) | hIRE1 LD + MPZ1N-2X (2:1) | hIRE1 LD + MPZ1N-2X (4:1) | hIRE1 LD + MPZ1N-2X-RD (2:1) | hIRE1 LD + MPZ1N-2X-RD ( 4:1) |
|---|---|---|---|---|---|---|---|
| **Best-fit values** | | | | | | | |
| Plateau | 0.8190 | 0.8138 | 0.9307 | 0.3492 | 0.6607 | 0.8577 | 0.7212 |
| Tau | 243.7 | 236.1 | 260.0 | 512.6 | 375.0 | 274.3 | 190.9 |
| Half time [sec] | 169.0 | 163.7 | 180.2 | 355.3 | 260.0 | 190.2 | 132.3 |
| **95% CI (profile likelihood)** | | | | | | | |
| Plateau | 0.7924 to 0.8502 | 0.7968 to 0.8325 | 0.9114 to 0.9519 | 0.3362 to 0.3640 | 0.6206 to 0.7130 | 0.8414 to 0.8755 | 0.7019 to 0.7433 |
| Tau | 221.3 to 270.6 | 222.2 to 251.7 | 245.6 to 276.0 | 479.5 to 550.4 | 327.5 to 437.6 | 261.1 to 288.8 | 173.5 to 211.4 |
| Half-time | 153.4 to 187.5 | 154.0 to 174.4 | 170.2 to 191.3 | 332.4 to 381.5 | 227.0 to 303.3 | 181.0 to 200.2 | 120.2 to 146.5 |
| **Goodness of Fit** | | | | | | | |
| R squared | 0.8296 | 0.9350 | 0.9411 | 0.9737 | 0.8298 | 0.9600 | 0.7907 |

**Kettel et al., Supp. Table 4.** linked to the data in **Fig. 3G**

Fits of the FRAP curves of **mCherry-IRE1 LD-10His** or **mCherry-IRE1 cLD-10His** in condensates formed in solution

| | LD | cLD |
|---|---|---|
| **Best-fit values** | | |
| Plateau | 0.9251 | 0.2753 |
| Tau | 281.1 | 406.1 |
| Half-time | 194.9 | 281.5 |
| **95% CI (profile likelihood)** | | |
| Plateau | 0.8842 to 0.9756 | 0.2635 to 0.2893 |
| Tau | 249.7 to 320.6 | 371.8 to 447.0 |
| Half-time | 173.1 to 222.2 | 257.7 to 309.9 |
| **Goodness of Fit** | | |
| R squared | 0.8022 | 0.9316 |
